# Ngfr suppresses Lcn2/Slc22a17 signaling, induces neurogenesis and reduces amyloid pathology in the hippocampus of APP/PS1dE9 mouse

**DOI:** 10.1101/2022.08.20.504608

**Authors:** Tohid Siddiqui, Mehmet Ilyas Cosacak, Stanislava Popova, Prabesh Bhattarai, Annie J. Lee, Yuhao Min, Xue Wang, Mariet Allen, Özkan İş, Zeynep Tansu Atasavum, Badri N. Vardarajan, Ismael Santa-Maria, Giuseppe Tosto, Richard Mayeux, Nilüfer Ertekin-Taner, Caghan Kizil

**Affiliations:** German Center for Neurodegenerative Diseases (DZNE) within Helmholtz Association; 01307, Dresden, Germany; Department of Neurology, Columbia University Irving Medical Center; New York, NY 10032, USA; The Gertrude H. Sergievsky Center, College of Physicians and Surgeons, Columbia University, 630 West 168th Street, New York, NY, 10032, USA; Taub Institute for Research on Alzheimer’s Disease and the Aging Brain, Columbia University Irving Medical Center; New York, NY 10032, USA; Department of Neuroscience, Mayo Clinic Florida, Jacksonville, Florida 32224, USA; Department of Quantitative Health Sciences, Mayo Clinic Florida, Jacksonville, Florida 32224, USA; Department of Pathology and Cell Biology, Columbia University Irving Medical Center; New York, NY 10032, USA; Department of Psychiatry, College of Physicians and Surgeons, Columbia University, 1051 Riverside Drive, New York, NY, 10032, USA; Department of Neurology, Mayo Clinic Florida, Jacksonville, Florida 32224, USA

**Keywords:** Mouse, zebrafish, human, Alzheimer’s disease, neurogenesis, nerve growth factor receptor, Lipocalin 2, transcriptomics, proteomics, 3D human astrocyte culture, post-mortem human brain, amyloid, Tau, weighted gene co-expression network analysis, cell intrinsic differential gene expression

## Abstract

Neurogenesis relates to the brain resilience and is reduced in Alzheimer’s disease (AD). Restoring healthy levels of neurogenesis could have beneficial effects for coping with neurodegeneration. However, molecular mechanisms that could enhance neurogenesis from astroglial progenitors in AD pathology are largely unknown. We used lentiviruses to express *Ngfr* in the hippocampus of the APP/PS1dE9 mouse model of AD, histologically analyzed the changes in proliferation of neural stem cells and neurogenesis; performed single-cell transcriptomics, spatial proteomics, and functional knockdown studies. We found that expression of *Ngfr*, a neurogenic determinant in pathology-induced neuroregeneration in zebrafish, stimulated proliferative and neurogenic outcome in the APP/PS1dE9 AD mouse model. Ngfr suppressed reactive astrocyte marker Lipocalin-2 (Lcn2) in astroglia. Blockage of Lcn2 receptor, Slc22a17, recapitulated the neurogenic effects of NGFR, and long-term Ngfr expression reduced amyloid plaques and Tau phosphorylation. Furthermore, immunostaining on postmortem human hippocampi with AD or primary age-related Tauopathy and 3D human astroglial cultures showed that elevated LCN2 levels correlate with gliosis. By comparing transcriptional changes in mouse hippocampus, zebrafish brain, and human AD brains in terms of cell intrinsic differential gene expression analyses as well as weighted gene co-expression network analysis, we observed common potential downstream effectors of NGFR signaling, *C4B* and *PFKP*, that are relevant to AD. Our study links the regulatory role of an autocrine molecular mechanism in astroglia to the neurogenic ability and modulatory effects on amyloid and tau phosphorylation, opening new research avenues and suggesting that neurogenesis-oriented therapeutic approaches could be a potential clinical intervention for AD.

## Background

Generation of new neurons in adulthood reduces with vertebrate phylogeny ^1^. Neurogenic regions are spatially restricted in mammals, which limits the addition of new neurons to the existing circuitry. Generation of new neurons and continuous integration could contribute to the resilience and cognitive reserve of the brain ^2,3^, which could affect late-onset neurodegeneration. Recent studies showed declining neurogenesis in Alzheimer’s disease (AD) patients ^4,5^. Although not clear whether reduced neurogenesis could be one of the culprits of AD, therapies aiming at enhancing neurogenic output hold promise for reverting the age-related neurodegenerative conditions. However, the molecular mechanisms underlying neurogenesis are largely unknown.

The heterogeneity of the neurogenic ability across vertebrates is vast. Adult teleost fish such as zebrafish has a widespread regenerative ability, which is unparalleled in the vertebrate clades ^6,7^. Astroglia acting as neural stem/progenitor cells are one of the key cell types for neuronal regeneration ^8^. We previously generated an adult AD zebrafish model and found that a complex set of cellular crosstalk leads to production of more neurons despite the disease ongoing pathology ^9–12^. This model has strong parallelism to human AD brains in terms of molecular programs affected by AD pathology ^13^, served as a pre-clinical tool for drug screening ^14–16^ and helped identify genes associated with AD in humans ^17,18^. The extraordinary regenerative ability of astroglia in zebrafish in AD-like scenarios rely on set of molecular mechanisms, one of which is the pro-neurogenic activity through the expression of nerve growth factor receptor (*ngfra*) ^9,19^. *Ngfr* is not detected in mouse hippocampal astroglia, and we hypothesized that ectopic expression of *Ngfr* could alter the neurogenic properties and AD pathology in mouse brains. In this study, we show the effects of Ngfr signalling on enhancing neurogenesis and reducing AD pathology, when activated in the hippocampus of an AD mouse model. We also provide a mechanistic link between NGFR activity and downstream transcriptional regulation and phenotypic outcomes by using in vitro 3D human neurogenesis assay, large human AD cohorts, and cell intrinsic differential gene expression analyses. Our results point towards the relevance and importance of neurogenic molecular programs in modulating the pathogenesis of Alzheimer’s disease.

## Results

We generated lentiviral constructs that code for mCherry (control, Lv13) and mCherry and mouse *Ngfr* (Lv16) to express *Ngfr* /*p75NTR* in the astrocytes of the subgranular zone (SGZ) of mouse dentate gyrus (DG) (Fig. S1). These viral particles have enhanced efficiency for targeting astroglia as documented before ^20^, yet can transduce other cell types. Microinjection into mouse SGZ (Fig. 1A) results in targeting and transduction of DG (Fig. 1B). In control DG, Ngfr is expressed at the periphery of the DG but not in SGZ, while Lv16 transduction leads to ectopic expression (Fig. 1B). The expressed receptor is functional as one of the Ngfr ligands - BDNF - increases the proliferation of mouse DG astrocytes only when Ngfr is expressed (Fig. S1C, S1D). To determine the molecular programs that are altered after *Ngfr* expression in astrocytes, we performed single-cell transcriptomics by dissecting the DG from Lv13 and Lv16-transduced wild-type mouse brains (Fig. 1C, 1D, Fig. S1E). Clustering the cell types and plotting Lv16-dependent expression of Ngfr-mCherry (adding the fusion transcript as a synthetic read template that determined the expression of the transgene, Fig. 1C-E) revealed that Lv16 transduction can target astrocytes as well as microglia. and that astroglia is the highest targeted cell type (Fig. 1D). Control transductions with a control virus (Lv13) expressing only mCherry did not show any *Ngfr*-mCherry fusion construct expression (Fig. 1D). To determine the effects of *Ngfr* transduction on astrocytes, we performed differentially expressed gene (DEG) (Data S1) and pathway analyses (Data S2 and Data S3) within the astroglial cluster (Cluster 4) and showed that *Ngfr* expression reduced pathways related to neurodegenerative diseases, including AD, while induced pathways related to sustained homeostasis and growth, such as proteasome, mTOR signaling and cell cycle (Fig. 1F). Analyses of DEGs in astroglia showed a general reduction in astrocytic markers that are associated with reactive or non-neurogenic states (e.g.: *Gfap, Apoe, Hopx, Ndrg2, Aldoc, Id3*) and an overexpression of genes associated with proliferation and differentiation (e.g.: *Prdm2, Enc1*, *Egfr, Mcm7, Cdk13*) (Fig. 1G-I). This suggested that Ngfr signaling could promote proliferation and neurogenesis in mouse DG astroglia.

**Fig. 1.**
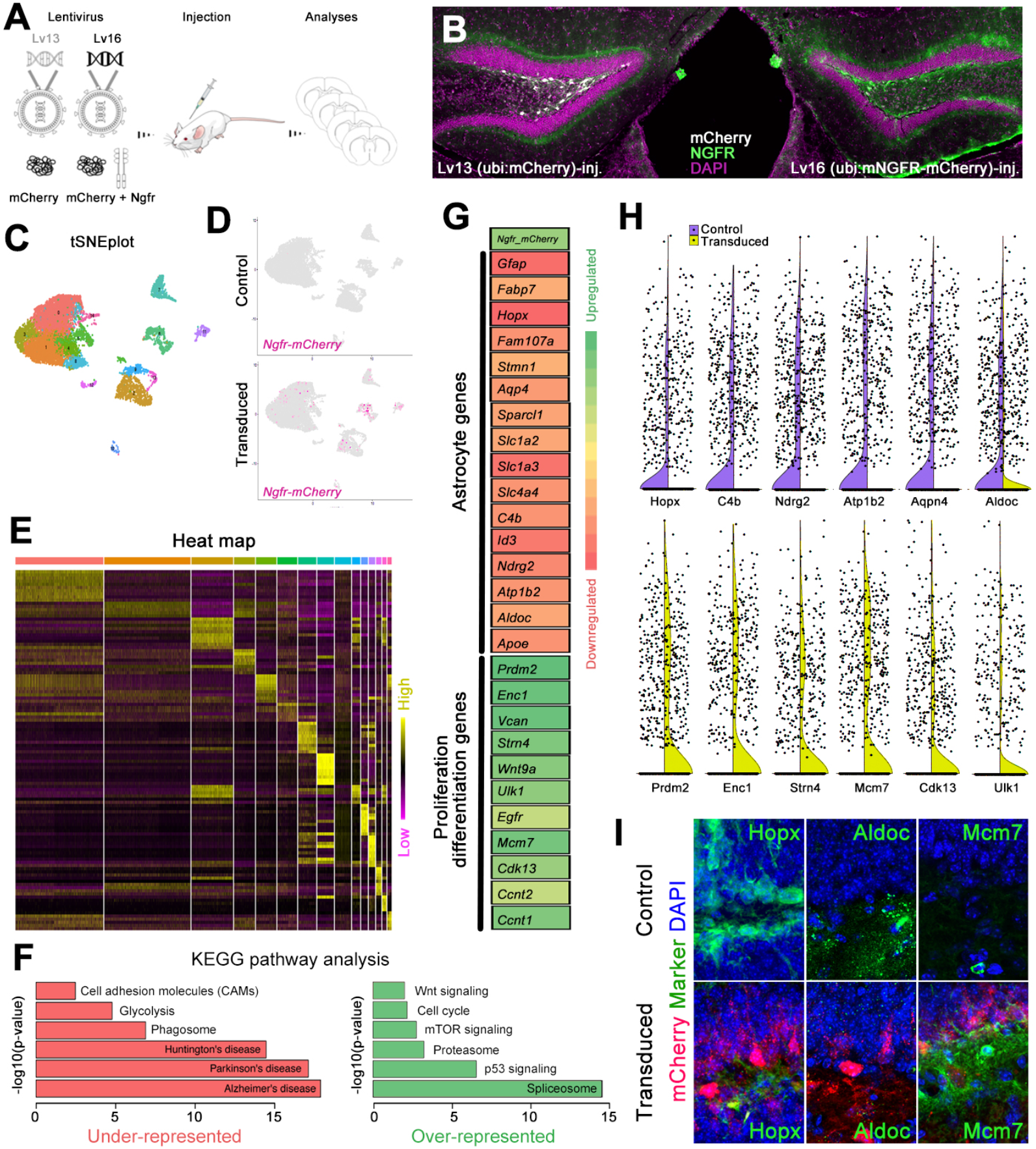
NGFR enhances proliferative and neurogenic markers in dentate gyrus (DG) astrocytes. **(A)** Schematic strategy for expression lentiviruses: Lv13: mCherry control; Lv16: Ngfr-mCherry. **(B)** Cross section images for DGs transduced with Lv13 and Lv16 and immunostained for mCherry, NGFR with DAPI counterstain. Note NGFR expression in the subgranular zone (SGZ) after Lv16 transduction. **(C)** Single cell transcriptomics tSNE plot from dissected DGs after Lv13 and Lv16 transduction. **(D)** tSNE plots from Lv13 and Lv16 transduction showing the expression of Ngfr-mCherry, which is detectable only after Lv16. **(E)** heat map of different cell types. **(F)** KEGG pathway analyses on astrocytic population, showing downregulated and upregulated pathways. **(G)** A heat map for selected differentially expressed genes (DEGs) after Lv16. Astrocyte markers are downregulated, and proliferation/neurogenesis markers are upregulated. **(H)** Violin plots for selected genes: purple: control (Lv13); yellow (Lv16). **(I)** Immunostaining for validating the DEGs in control and Lv16-transduced DG. Scale bars equal 50 μm.

To determine whether *Ngfr* expression could enhance proliferation and neurogenesis from astroglia in DG, we performed Lv13 and Lv16 transduction, BrdU labeling, and quantified the extent of labelled astroglia (Gfap+) and newborn neurons (Dcx+) at 3 days after transduction in wild type and APP/PS1dE9 mouse model of AD (Fig. 2, Fig. 3, and Fig. S2). We found that Ngfr expression enhanced proliferation and neurogenesis in both healthy (Fig. 2A-F, Fig. S2, and Fig. 3; Data S4; by 2.99-fold and 2.08-fold, respectively) and AD mouse brains (Fig. 2G-L, Fig. S2, and Fig 3; Data S4; by 4.85-fold and 7.32-fold), validating the Ngfr-mediated molecular changes in astrocytes (Fig. 1). To determine the effect of Ngfr on proliferation and neurogenesis in astrocytes, we studied the Ngfr-transduced astroglia in our single-cell sequencing by *in silico* dissection of Ngfr-mCherry expressing cells and determining the DEGs in comparison to non-transduced astroglia from the same mouse brain (analysis 1) or astroglia from control virus (Lv13) transduced mouse brain (analysis 2) (Fig. 4A, Data S2 and Data S3). We performed these analyses to ensure that Ngfr expression in astroglia is not a result of technical variables (by comparing the same astroglia population in the same animal according to their transduction status) or due to the presence of the lentiviral particles in the brain (comparing the astroglia from control Lv13 and Ngfr-containing Lv16-transduced animals). When we selected the top-50 DEGs and selected the common hits in analyses 1 and 2, we found 10 genes that changed their expression upon NGFR expression in both analyses, confirming the robustness of our findings (Fig. 4A). We hypothesized that if a specific cell autonomous effect of Ngfr is running through a select number of genes in neurogenic cell populations, then those genes could be mainly expressed in astroglia. Investigating the genes only expressed in astroglia would also minimize the effects of non-astroglial transduction of the viral particles. To determine whether any of the identified candidate genes has restricted astroglial expression, we generated tSNE plots and found that *Gm8251, Cxcl1*, and *Lcn2* were expressed predominantly in astroglia (Fig. 4B). By drawing Vln plots, we observed that only *Lcn2* showed a significant change in expression in astroglia (Fig. 4C). To validate our findings, we performed immunohistochemical staining for Lcn2 in Lv16-transduced DG and found that while non-transduced astroglia were Lcn2-positive, Ngfr-mCherry positive astroglia did not express Lcn2 (Fig. 5D), supporting the hypothesis that NGFR signalling specifically reduces Lcn2 in DG astroglia.

**Fig. 2.**
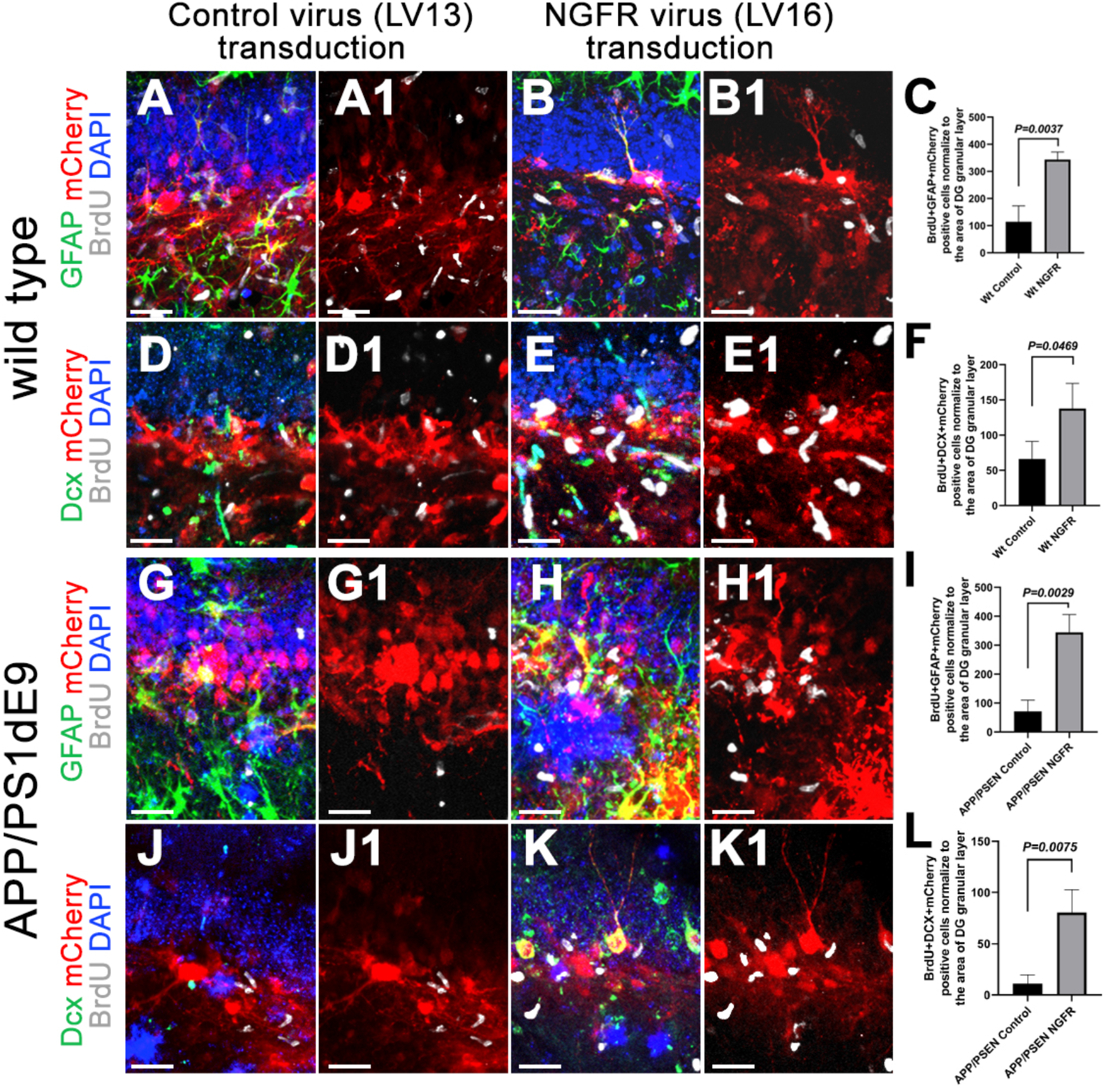
NGFR promotes proliferation of DG astrocytes and neurogenesis in wild type and APP/PS1dE9 model of AD. **(A, B)** Immunostaining for GFAP, BrdU and mCherry with DAPI counterstain in Lv13-(A) and Lv16-(B) SGZ of wild type mice. **(A1, B1)** BrdU/mCherry channels of A and B. **(C)** Quantification graph for mCherry-BrdU-GFAP triple positive cells. **(D, E)** Immunostaining for Dcx, BrdU and mCherry with DAPI counterstain in Lv13-(A) and Lv16-transduced (B) SGZ. **(D1, E1)** BrdU/mCherry channels of D and E. **(F)** Quantification graph for mCherry-BrdU-Dcx triple positive cells of wild type mice. **(G, H)** Immunostaining for GFAP, BrdU and mCherry with DAPI counterstain in Lv13-(G) and Lv16-transduced (H) SGZ of APP/PS1dE9 mice. **(G1, H1)** BrdU/mCherry channels of G and H. **(I)** Quantification graph for mCherry-BrdU-GFAP triple positive cells. **(J, K)** Immunostaining for Dcx, BrdU and mCherry with DAPI counterstain in Lv13-(J) and Lv16-transduced (K) SGZ of APP/PS1dE9 mice. **(J1, K1)** BrdU/mCherry channels of J and K. **(L)** Quantification graph for mCherry-BrdU-Dcx triple positive cells. Scale bars equal 50 um.

**Fig. 3.**
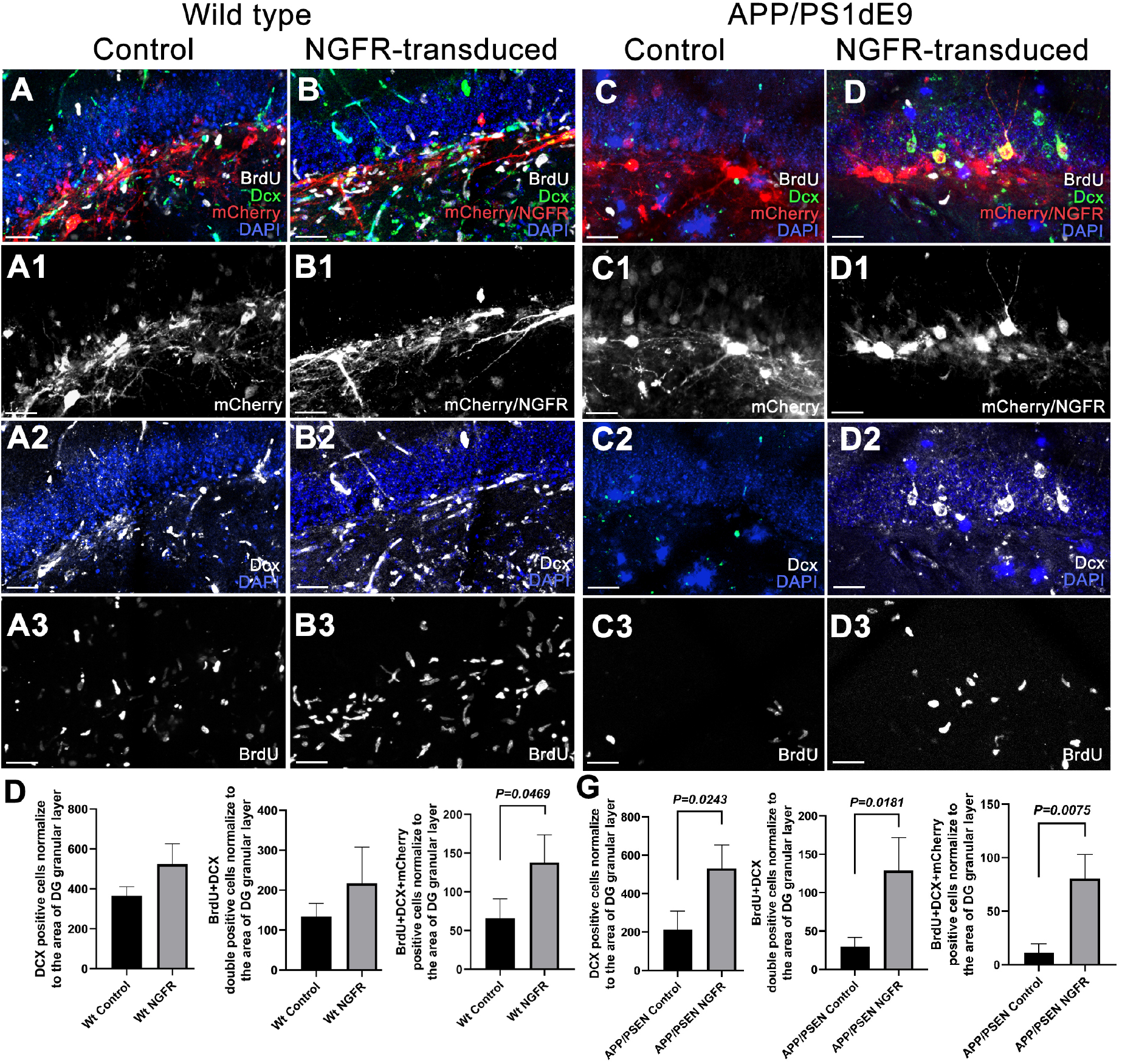
Lv16 transduction enhances neurogenesis in wild type and APP/PS1dE9 mouse model of Alzheimer’s disease. **(A)** Dcx, mCherry, and BrdU triple immunostaining with DAPI counterstain in Lv13-transduced wild type mouse dentate gyrus. **(A1)** mCherry channel of A. **(A2)** Dcx and DAPI channel of A. **(A3)** BrdU channel of A. **(B)** Dcx, mCherry, and BrdU triple immunostaining with DAPI counterstain in Lv16-transduced wild type mouse dentate gyrus. **(B1)** mCherry channel of B. **(B2)** Dcx and DAPI channel of B. **(B3)** BrdU channel of B. **(D)** Normalized values of Dcx-positive, BrdU-Dcx double positive and BrdU-Dcx-mCherry triple positive cells in wild type mouse dentate gyrus transduced with Lv13 (control) or Lv16 (NGFR). **(E)** Dcx, mCherry, BrdU triple immunostaining with DAPI counterstain in Lv13-transduced APP/PS1dE9 mouse dentate gyrus. **(E1)** mCherry channel of e. **(e2)** Dcx and DAPI channel of E. **(E3)** BrdU channel of E. **(F)** Dcx, mCherry, and BrdU triple immunostaining with DAPI counterstain in Lv16-transduced wild type mouse dentate gyrus. **(F1)** mCherry channel of F. **(F2)** Dcx and DAPI channel of F. **(f3)** BrdU channel of F. **(G)** Normalized values of Dcx-positive, BrdU-Dcx double positive and BrdU-Dcx-mCherry triple positive cells in APP/PS1dE9 mouse dentate gyrus transduced with Lv13 (control) or Lv16 (NGFR). Scale bars equal 50 μm.

**Fig. 4.**
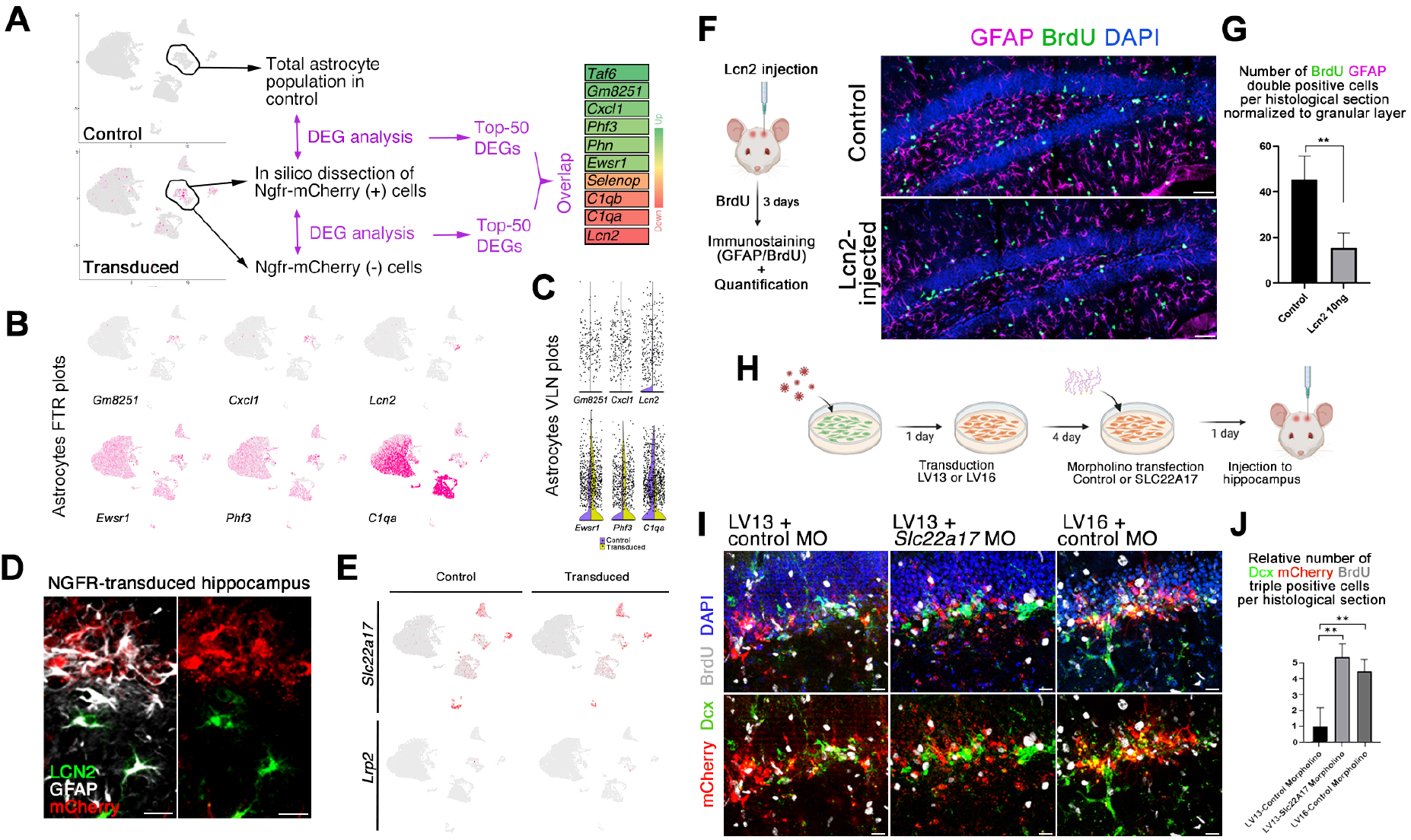
Ngfr regulates neurogenic response in DG astrocytes through suppression of Lcn2/Slc22a17 activity. **(A)** Single cell transcriptomics strategy for isolating Ngfr-mCherry positive astrocytes and comparing the transcriptomics profiles to non-transduced astrocytes or control astrocytes. The comparison showed differential expression of 10 genes in common as shown in the heat map. **(B)** Expression of selected genes on tSNE plots. **(C)** Violin plots for selected genes. **(D)** Immunostaining for Lcn2, GFAP and mCherry in LV16 Ngfr-transduced DG. Lcn2-positive and mCherry positive astrocytes do not overlap. **(E)** tSNE plots for two receptors of Lcn2: *Lrp2, Slc22a1**7***. **(F)** Injection, BrdU treatment and analyses scheme of the effect of Lcn2 on proliferating GFAP cells, and immunohistochemical staining for BrdU, GFAP and DAPI. **(G)** Quantification graph for BrdU-GFAP double positive cells. **(H)** Schematic representation of the cell-type specific functional knockdown of *Slc22a17* in astrocytes. **(I)** mCherry, Dcx, BrdU immunostaining with DAPI counterstain in SGZs transplanted with Lv13+control morpholino, Lv13+*Slc22a17* morpholino and Lv16+control morpholino-treated astrocytes. Lower panels: DAPI omitted from upper panels **(J)** Quantification of Dcx, mCherry and BrdU triple positive cells in i. Slc22a17 knockdown mimics Ngfr transduction. n = 3. Scale bars equal 25 μm.

**Fig. 5.**
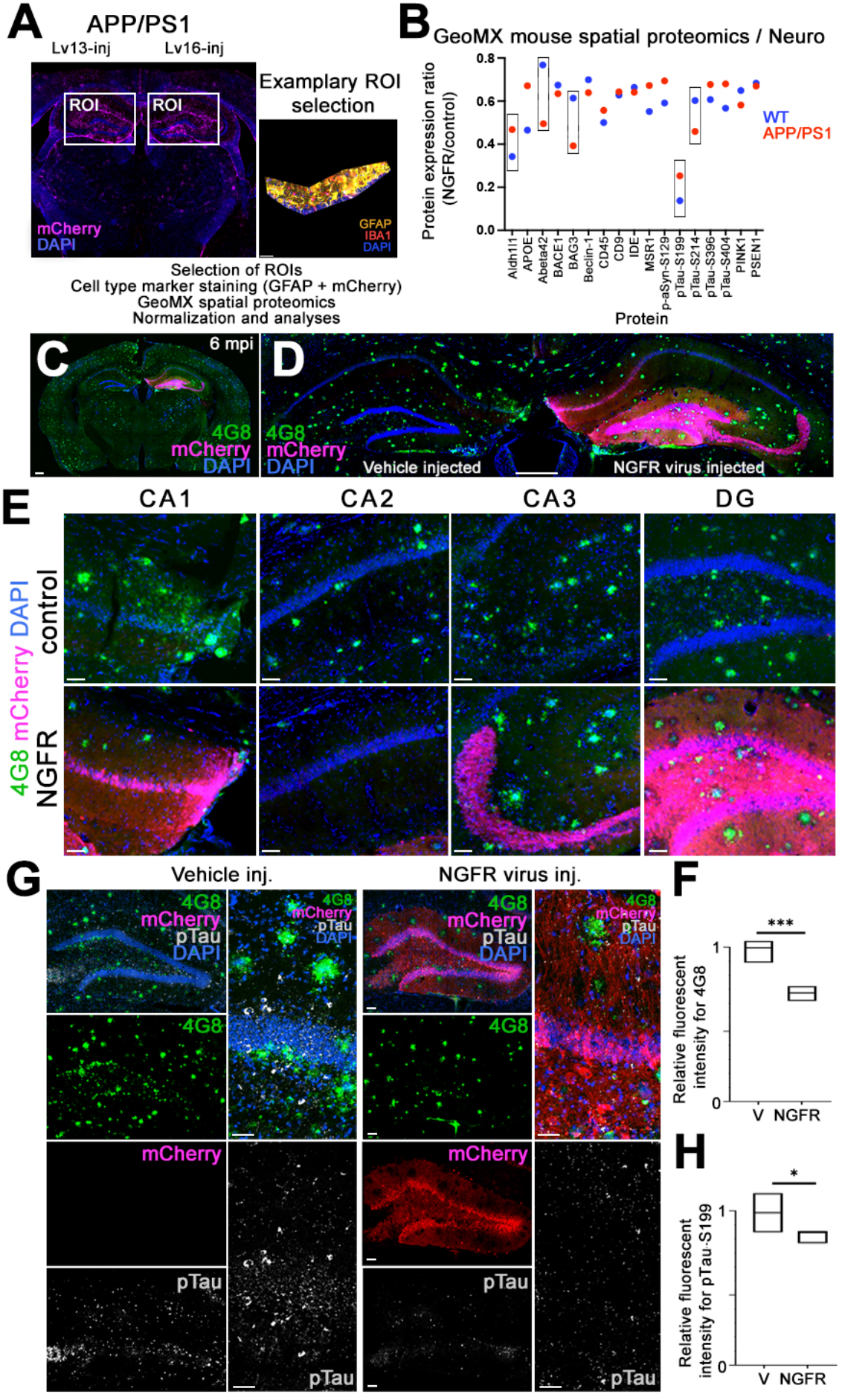
Ngfr reduces amyloid-beta42 load and phosphorylated Tau in the hippocampus of APP/PS1dE9 mice. **(A)** Representative image from GeoMx spatial proteomics mouse brain section and region of interest (ROI) at the SGZ. **(B)** Plot showing the fold changes of selected proteins upon *Ngfr* transduction in wild type and APP/PS1dE9 mice. **(C)** Representative APP/PS1dE9 mouse brain section immunostained for 4G8 and mCherry with DAPI counterstain at 6 months after transduction with Lv16 in one DG. **(D)** High-magnification image from C. **(E)** Comparison of amyloid load in CA1, CA2, CA3 and DG regions of control and *Ngfr*-transduced hemispheres. **(F)** Quantification of amyloid load in terms of normalized amyloid immunoreactivity by comparing control (v) versus *Ngfr* (Lv16) transduction. n = 5. **(G)** 4G8, mCherry, phosphorylated Tau-S199 immunostaining with DAPI counterstain in control and *Ngfr*-injected DGs. Individual fluorescence channels and close-up images are also shown. **(H)** Quantification of pTau-S199 in terms of normalized fluorescence by comparing control (v) versus *Ngfr* (Lv16) transduction. n = 5. ***: p<0.005, *: p<0.05. Scale bars equal 100 μm.

Lcn2 is a ligand and can bind to two receptors, Lrp2 and Slc22a17 ^21,22^. To determine whether these receptors are expressed in mouse DG, we generated two tSNE plots for the receptors and found that, while *Lrp2* is not expressed, *Slc22a17* is expressed in several cell types, including astroglia (Fig. 4E), suggesting that Lcn2 can act via an autocrine signalling. To determine whether Lcn2 could alter the proliferation of astroglia, we injected Lcn2 into the hippocampus of the wild-type mice, treated the animals with BrdU and analyzed the labelled astroglia (Fig. 4F). We observed that Lcn2 significantly reduces the number of label-retaining Gfap+ cells, indicating reduced astroglial proliferation (Fig. 4G). To determine whether Lcn2-Slc22a17 signalling is biologically relevant to neurogenesis, we performed loss-of-function studies for *Slc22a17* (Fig. 4H-J) by using morpholino oligonucleotides that effectively reduce Slc22a17 protein levels (Fig. S3B-D). Since Slc22a17 is expressed in several cell types and to avoid pleiotropic effects of the loss-of-function that would confound the astroglia-related observations, we transduced the mouse DG astrocytes *in vitro*, treated the cells with *Slc22a17* morpholinos and then transplanted these astroglia to the SGZ of wild type mouse brains (Fig. 4H). With this method, specific blockage of Lcn2-Slc22a17 signalling in mCherry-positive astroglia could be achieved and assessed without altering this signalling pathway in other cell types. To determine the progeny of the transplanted astroglia, we injected the mice with BrdU after transplantation. After performing immunohistochemical staining for BrdU (newborn cells), Dcx (early neurons) and mCherry (transduced cells), we found that compared to Lv13 and control morpholino, Lv13 and *Slc22a17* morpholino significantly increased the generation of newborn neurons from transplanted astrocytes and this increase is comparable to Lv16 (Ngfr) transduction alone (Fig. 4I, J; Data S4). These results suggest that the imposition of neurogenic potential by Ngfr expression to DG astroglia might be cell autonomous through autocrine Lcn2-Slc22a17 signalling.

Since Ngfr signalling in DG astrocytes induced proliferation and neurogenesis (Fig. 2), and suppressed the molecular pathways related to AD (Fig. 1), we hypothesized that active Ngfr signalling could alter AD pathology ^4,23–26^. To determine whether Ngfr signalling would alter AD-related proteins that cannot be identified by transcriptomics, we performed spatial proteomics on mCherry-enriched regions of Lv13 and Lv16-transduced SGZs in wild type and APP/PS1dE9 AD model (Fig. 5A). After data normalization and quality control (Fig. S4, Data S7 and Data S8), we plotted the fold changes in protein expression levels between Ngfr and control transduction (Fig. 5B, Fig. S5 and Fig. S6). We found that the highest fold changes upon Ngfr transduction in APP/PS1dE9 mouse was observed for Aldh1l1 (−63% in wt, −53% in APP/PS1), amyloid beta −42 (−21% in wt, −47% in APP/PS1), phosphorylated Tau (S199) (−86% in wt and −75% in APP/PS1), phosphorylated Tau (S214) (−40% in wt, −54% in APP/PS1), and BAG3 (−39% in wt, −61% in APP/PS1) (Fig. 5B). Other AD-related proteins such as pTau-S396, pTau-S404, and APOE were also downregulated by NGFR (Fig. 5B, Data S8), suggesting that activation of NGFR signalling in DG and particularly in astroglia could improve AD pathology burden. To test this hypothesis, we transduced the APP/PS1dE9 mice with Lv16 and analyzed these brains 6 months after transduction (Fig. 5C). mCherry signal in the transduced hemisphere covered the entire DG, indicating that Ngfr-transduced astrocytes contributed to neurogenesis (Fig. 5D). Additionally, when different subregions of the hippocampus were analyzed, we observed reduced immunoreactivity for 4G8-positive amyloid plaques (Fig. 5E), and this reduction amounted to 22% in the entire hippocampus (Fig. 5F). Since spatial proteomics showed reduced levels of phosphorylated Tau (Fig. 5B), we tested these findings by performing immunohistochemical staining for pTau-S199 in control and Ngfr-transduced hemispheres of APP/PS1dE9 mouse DG. We observed that Ngfr-transduced DGs have significantly lower amounts of pTau-S199 in the overall hippocampus (11%, Fig. 5G, H), while the difference in SGZ is more pronounced (Fig. 5G). Overall, these results suggest that Ngfr signalling could reduce amyloid and Tau pathology by enhancing neurogenesis in mouse hippocampus.

To determine whether Ngfr receptor expression in human brains would correlate with proliferating cells, we analyzed three publicly available single-cell sequencing datasets (human brain organoids, fetal human brain, adult human brain) (Fig. S6) ^27,28^. Cell clustering, marker gene identification, and cell type determination followed by generation of tSNE plots for *NGFR* and *PCNA* (proliferating cell marker) showed that in brain organoids and developing fetal brain, *NGFR* expression partially overlaps to proliferating astroglial cells while in adult human brains, *NGFR* is not expressed in astroglia (Fig. S6), suggesting an age-related loss of *NGFR* activity and corresponding reduced neurogenesis.

To correlate our findings to human brains, we hypothesized that the expression of *LCN2* could increase in the brains of AD patients, where neurogenesis is reduced. To test this hypothesis, we performed immunolabeling of human hippocampi for GFAP and LCN2 in healthy controls and AD patients (Fig. 6). We observed scarce LCN2 expression in healthy aged individuals (Fig. 6A, B), while in human AD brains LCN2-positive astroglia increases dramatically concomitant to architectural changes of glial extensions and elevated hypertrophy (Fig. 6C, D). Quantification of LCN2-positive GFAP+ cells normalized to the overall GFAP+ cells indicate that the number of LCN2-positive glia increases significantly in late-stage AD patients (Braak stage V-VI) but not in earlier AD stages (Braak stage IV-V) (Fig. 6E). We observed that the LCN2-expressing astroglia are more compact than their healthy counterparts (Fig. 6B, D). To determine whether LCN2 is sufficient to alter the morphology of astroglia, we used 3D hydrogel cultures of primary human astroglia (Fig. 6F). We observed that treatment of astroglia with LCN2 significantly reduces the glial volumes indicative of a reactive state (Fig. 6G), consistent with the reduction of GFAP and MAP2-positive cells, indicative of reduced neurogenic potential (Fig. 6H, I). To further investigate the role of NGFR signalling in AD, we compared the transcriptional changes exerted by *Ngfr* expression in DG of APP/PS1dE9 mouse (Data S6) to large AD human cohorts in humans (Fig. 7). We used the ROSMAP study and its bulk RNA sequencing from the dorsolateral prefrontal cortex (DLPFC), anterior caudate (AC) and posterior cingulate cortex (PCC) regions ^29,30^ (Fig. 7A). To add another stringency level and a biologically relevant experimental animal model, we also included data from sorted astroglia of an amyloid toxicity model of adult zebrafish brain, where the induced pathology elicits an Ngfr-dependent neurogenic response ^9,13^ (Fig. 7A). In mouse astroglia transduced with *Ngfr* vs. control astroglia, we identified 152 genes that are significantly DEGs, of which 61 are also nominally significant DEG in at least one ROSMAP brain region (Fig. 7B, Data S9). We hypothesized that the pathology-altering and neurogenic effects of Ngfr activity could be reflected in the gene expression patterns of these overlapping 61 genes. Hypothetically, in mouse and human AD brains, the genes must have opposite directionality of change of expression. We found 15 genes conform to this criterion (Fig. 7B, Data S6). Since zebrafish astroglia becomes neurogenic after amyloid toxicity upon *Ngfr* signalling, we used zebrafish brain as a reference comparison and hypothesized that the expression change in those genes should be same direction with mouse but opposite direction with the human AD cohorts. 7 genes conformed to these criteria (Fig. 7B, C). Their expression heatmap indicated that in human AD brains, *WDR53, GADD45B, and GLN3* are downregulated while *C4B, PFKP, S100A6* and *SELENOP* are upregulated (Fig. 7C), while *Ngfr* transduction in mouse hippocampus or amyloid toxicity in adult zebrafish brain changed these genes in opposite directionality (Fig. 7C).

**Fig.6.**
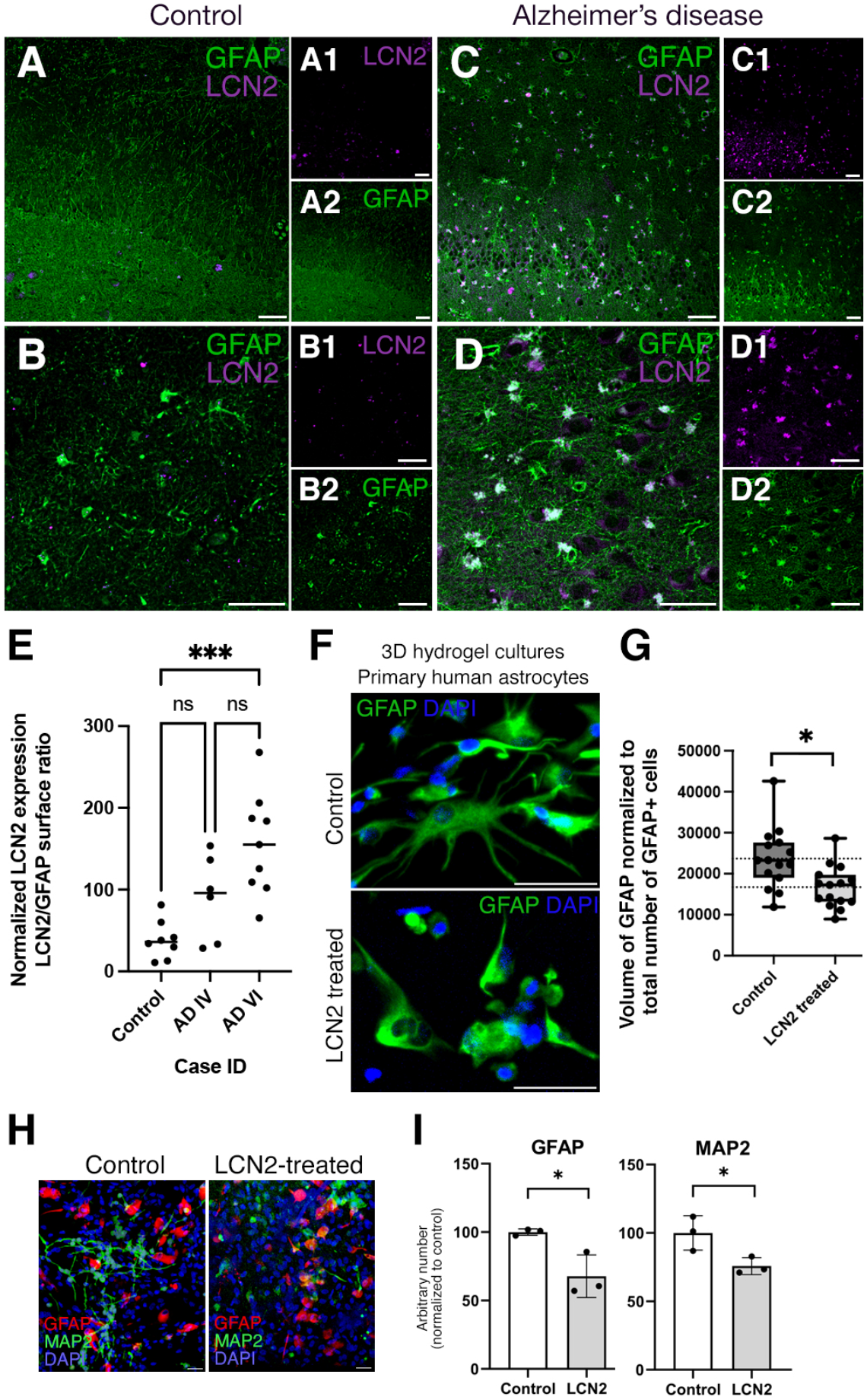
LCN2 is upregulated in human brains with AD. **(A-D)** Immunohistochemical stainings for LCN2 and GFAP on hippocampal brain sections of healthy control (A, B) and Braak stage VI AD patient (C, D). Black-white insets (A1-D2) indicate individual fluorescent channels. LCN2 (top) and GFAP (bottom). **(E)** Quantification of LCN2-positive GFAP cells normalized to total GFAP cells. In total n = 7 human brains, n = 21 images analyzed. ***: p<0.001. **(F)** Immunostaining for GFAP on primary human astrocytes in 3D hydrogels: control and LCN2-treated. **(G)** Quantification of the volume of GFAP normalized to total number of GFAP cells. LCN2-treatment reduces the volume of astroglia, indicative or reactive states. Scale bars equal 50 μm. **(H)** GFAP and MAP2 immunostaining on control and LCN2-treated 3D hydrogel cultures of human primary fetal astroglia. **(I)** Quantification of normalized GFAP and MAP2-positive cells after the culture period.

**Fig.7.**
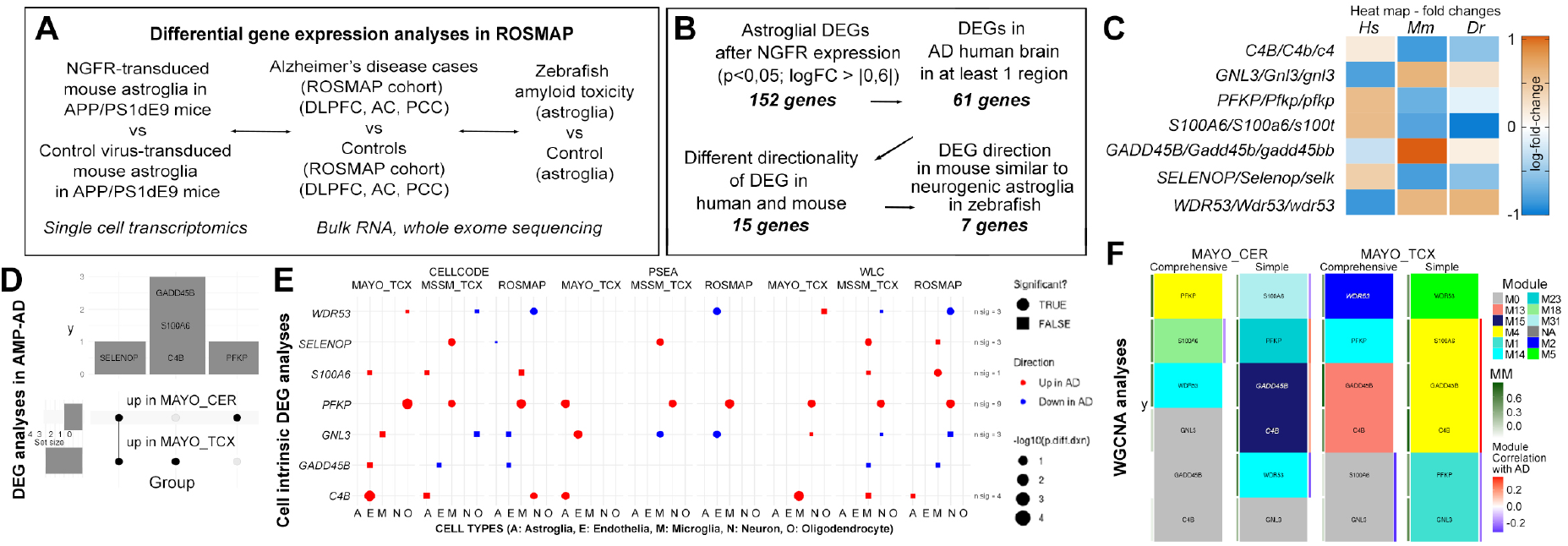
Comparison of gene expression changes in mouse brain with NGFR to human AD cohorts by differential gene expression analyses, cell intrinsic gene expression and Weighted gene co-expression network analysis. **(A)** Schematic flow of the differential gene expression analyses with ROSMAP AD cohort. **(B)** Stringency criteria and number of genes in each comparison category. **(C)** Heat map of expression changes of 7 candidate genes. Hs: human, Mm: mouse, Dr: zebrafish. **(D)** Overlap of significant DEG (FDR < 0.05 in the AMP-AD datasets. **(E)** Cell-intrinsic DEGs in 5 different cell types calculated using 3 different analytic tools (CellCODE, PSEA, WLC) from 3 different datasets (Mayo, MSSM, ROSMAP). Color indicates direction of changes. Circles are significant changes (p<0.05) while squares are not. **(F)** The gene of interest and their assigned modules (tile color) in WGCNA networks constructed from Mayo CER and TCX datasets either adjusting (comprehensive) or not adjusted (simple) for the cell proportion changes. Red-Blue color tile to the right indicates module correlation to AD diagnosis, where only significant correlation (p < 0.05) is shown. Green tile to the left indicates the gene’s module membership with respect to its assigned module.

To further analyze the involvement of these 7 genes in AD brains, we assessed their expression by three different approaches. At the bulk tissue level, we collected DEGs comparing AD and control brains from the AMP-AD datasets representing seven distinct datasets ^29,31–33^ (Fig. 7D). 2) For bulk analyses, only those genes differentially expressed at FDR<0.05 were considered DEGs. At the cell type level, we retrieved the cell-intrinsic DEGs (CI-DEGs) comparing gene expression levels between AD and control brains obtained using deconvoluted bulk gene expression from three independent bulk gene expression datasets via three different methods as previously described ^34^ (Fig. 7E). 3) At the gene network level, we retrieved the information about the co-expression modules ^35^ where the gene of interest is a member of the network module. The modules were constructed using AMP-AD Mayo Clinic temporal cortex (Mayo TCX) and cerebellum (CER) datasets as previously described ^36,37^ (Fig. 7F). Two analytic models were applied, a comprehensive model that adjusts for cell type markers to account for cell population variation in bulk tissue, and a simple model that does not. Relevant network information includes the module membership, module correlation with AD diagnosis, as well as enriched GO terms in the modules (Figure 7D-F, Data S10).

Five of the 7 genes are DEGs (FDR < 0.05) in the Mayo Clinic AD cohorts analyzed, although nominally significant DEGs were observed in other cohorts for these and other genes (Data S10). *SELENOP* is up in both Mayo CER and TCX, *C4B*, *S100A6*, and *GADD45B* are up in Mayo TCX, while *PFKP* is up in Mayo CER (Fig. 7D, Data S10). In the CI-DEG datasets, 6 out of the 7 genes were significantly (p <0.05) differentially expressed in at least one of the cell types in any of the three brain regions (Fig. 7E, Data S10). *PFKP* showed consistent upregulation in AD. Its upregulation is detected in neurons in 4/9 analyses, microglia (N=3), astrocytes (N=1), and oligodendrocytes (N=1), but not in endothelia. Another gene that has CI-DEGs in astrocytes (N=3) is *C4B*, which is also up-regulated in endothelia (N=1), microglia (N=2), and neurons (N=1). These findings provide further support for human brain expression changes in *C4B, PFKP, S100A6* and *SELENOP* that are biologically congruent with those from mouse and zebrafish models. Of these *C4B* and *PFKP* are also consistent CI-DEGs in neurons and glia.

Of the 4 types of co-expression modules evaluated (2 brain regions, 2 analytic models), *C4B* resides within Mayo TCX Simple Module M4, TCX Comprehensive M13, and CER Simple M15 (Fig. 7F, Data S10). These modules are enriched in microglia and endothelia genes, where *C4B* is a highly connected hub gene in the TCX Simple M4 (module membership=MM=0.8). Further both CER Simple M15 and TCX Simple M4 have positive correlations with AD diagnosis, consistent with the DEG and CI-DEG results. Interestingly, *GADD45B* resides in the same modules as *C4B*, suggesting co-regulation of these genes.

*PFKP* is a member of the TCX Simple M1, TCX Comprehensive M14, and CER Comprehensive M4 modules, all of which are enriched for neuronal genes, in addition to CER Simple M23. *PFKP* is a hub gene for both CER but not for the TCX modules. Both Simple modules CER M23 and TCX M1 are correlated with AD, former showing higher levels, consistent with DEG results, though latter showing lower levels, likely driven by other genes in this neuronal module enriched for synaptic biological terms.

Taken together, the human brain gene expression data supports robust expression changes in AD brains in a direction congruent with the cross-species data, and evidence of neuronal and glial expression perturbations for *C4B* and *PFKP* that are co-expressed with other microglial/endothelial and neuronal genes, respectively.

## Discussion

The reduced neurogenesis outcome in Alzheimer’s disease could be a pathological culprit of the disease ^4,5,23^. Therefore, enhancing neurogeniesis in neurodegenerative disease conditions could help to restore the brain’s resilience and ability to cope with neurodegeneration ^24,26^. However, the understanding of the molecular mechanisms of such a neurogenic competency is insufficient. Previous studies showed that neurotrophin signalling, especially BDNF, imposes a neuroprotective ability ^38,39^ and can contribute to the amelioration of cognitive decline in AD ^25,40^. Major neurotrophin receptor TrkB was associated with neuronal survival; however, BDNF is not directly inducing neural progenitor proliferation and neurogenesis in rodent brains ^25^. We previously showed that BDNF signalling imposed neuroregenerative ability through direct induction of astroglial proliferation and neurogenesis in an experimental adult zebrafish AD brain model ^9^. We found that this activity is dependent on NGFR/p75NTR receptor but not TrkB. NGFR can be bound by various ligands such as BDNF or NGF or their pro forms ^41^. In the hippocampus, we did not detect active cleaved forms of BDNF or NGF in the SGZ, but pro-BDNF is expressed in this region (Fig. S7). This is consistent with the binding ability of pro-BDNF to NGFR ^42^, and suggests that NGFR/pro-BDNF signalling could be responsible for neurogenic activity of mouse astrocytes. Therefore, here we show that transferring the molecular understanding from organisms with neuroregenerative ability, such as zebrafish, to mammalian brains is a promising cross-species approach to study the neuroregenerative ability to counteract AD pathology.

We found that *Lcn2* expression is negatively regulated by Ngfr signalling in mouse astroglia, and *Lcn2* and its receptor *Slc22a17* negatively regulates astrocyte proliferation and neurogenesis. This ultimately means that neurogenesis can be modified by targeting specific signalling pathways and the resilience of the brain in disease can be modified. Lcn2 is a secreted protein that is associated with autocrine induction of reactive astrocyte state and induce hypertrophic Gfap expression ^43^. *Lcn2* is upregulated in several brain diseases and may contribute to neurodegeneration, promote cell death and inflammation ^44^. In this study, we found that LCN2 is also upregulated in human AD, consistent with previous findings ^45^. However, the link between Ngfr signalling in astrocytes and Lcn2 has not yet been documented. Here, we propose that Ngfr signalling downregulates Lcn2 and promotes neurogenesis by altering the molecular signatures of reactive astrogliosis. Our spatial proteomics and single-cell transcriptomics data support this hypothesis (Fig. S7, Data S6). Additionally, our spatial proteomics analyses showed that, among the immune-related proteins, Cd44 was the most downregulated gene after Ngfr expression (Fig. S5), which is consistent with previous independent findings that LCN2 promoted CD44 ^46^.

We propose that NGFR signalling is an evolutionary determinant of the neurogenic potential in astroglia. In zebrafish, Ngfr signalling is naturally active in neural progenitor cells throughout the lifespan, which correlates with the life-long proliferation and neurogenic activity ^9^, while in the mouse brain, Ngfr is not present in hippocampal astroglia. When Ngfr is activated, reactive astrocyte programs are suppressed through suppression of Lcn2 and neurogenic programs prevail. Therefore, we hypothesize that mammalian brains could have lost NGFR expression in adult brain astroglia through the evolution, and this could have restricted the neurogenic ability in healthy and diseased mammalian brains. This hypothesis is supported by findings that a small population of astrocytes (0.3%) in the subventricular zone (SVZ) of mouse astrocytes are Ngfr-positive and they constitute a highly proliferative and neurogenic subset ^47^, and Ngfr signalling could regulate genes related to cell proliferation, differentiation and neurite growth *in vitro* ^48^. Our findings on single-cell sequencing data of human brain organoids, fetal human brain, and adult human brains (Fig. S6) also confirmed that *NGFR* expression in astroglia declines with age and could contribute to the hampered neurogenic outcome or inability of adult human brains to fulfil efficient neuroregeneration. In AD patients, LCN2-positive astroglia increase (Fig. 6), suggesting that NGFR/LCN2/SLC22A17 signaling axis could be a critical fate determination step between neurogenic versus reactive gliotic response in disease. In primary age-related Tauopathy, LCN2 expression does not change significantly in astroglia (Fig. S7), which supports the findings that amyloid toxicity could be the cause of LCN2 altered expression and subsequent AD pathology, including Tau phosphorylation. pTAU-S199 is a critical residue that is phosphorylated by CDK5 at late stages before the neurofibrillary tangle formation and is one of the residues detected by AT8 immunostaining ^49^. The reduction in Amyloid beta42 and phosphorylated forms of Tau protein could explain the long-term benefit of active NGFR signalling in astroglia.

We identified a small set of genes that might be regulated by NGFR signalling in vertebrate brains (Fig. 7, Data. S9, Data. S10). Particularly, *PFKP* and *C4B* are upregulated in human AD cohorts (Fig. 7) but downregulated with *Ngfr* transduction in astroglia as well as upon amyloid toxicity in adult zebrafish brain (Fig. 7). *C4B* is a complement protein that is involved in classical activation pathway ^50^, and found in AD patients cerebrospinal fluid as a biomarker for disease progression ^51^. Our analyses found *C4B* in human brains upregulated in bulk tissue, astrocytes and other glia and its expression correlates with the expression modules enriched for microglial/endothelial genes (Fig. 7, Data S10). This suggests that the expression of *C4B* in astroglia could have an immunomodulatory effect regulated by NGFR signalling. This hypothesis is reasonable, as astroglia can modulate the immune environment and the progression of disease pathology ^18,52–54^. *PFKP* is an enzyme that regulates the glycolytic pathway. We found this gene consistently upregulated in human AD brains (Fig. 7, Data S9, Data S10) and residing in neuronally enriched expression modules correlated with AD (Fig. 7F, Data S10). Alteration of energy metabolism in AD is a prominent pathological mechanism ^55,56^ and regulation of *PFKP* by NGFR signalling could be a mechanistic link to AD pathology. Further investigations on these two and other candidate genes could link the neurogenic outcome to the amelioration of the AD pathology in mammalian brains based on our mouse and zebrafish experiments and human AD cohort results.

It is unclear how the neurogenic effects of NFGR, anti-reactive gliotic role for the suppression of *Lcn2* expression, and other NGFR-mediated mechanisms act in concert to alleviate the pathological hallmarks of AD in long-term, yet this is an intriguing and new research avenue. Our single cell transcriptomics data comparing Ngfr-transduced brains versus controls showed that in neuronal, microglial and oligodendrocyte cell populations, molecular pathways related to unfolded protein response, clearance of misfolded proteins, protein processing, proteasome and Tau protein binding were enriched (Fig. S8). This suggests that NGFR signalling may induce more efficient clearance of toxic proteins, which might explain the long-term reduction of amyloid and Tau burden in Ngfr-transduced mouse brains (Fig. 5). This is consistent with previous findings ^57^. Recently, we showed that astroglial end-feet need to be retracted from the blood vessels to enhance efficient clearance of amyloid aggregates, and genetic mutations in humans alter this ability ^18^. While astroglia undergo proliferation, they retract their end-feet ^57–61^ and this could help enhance clearance. Longitudinal studies in animal AD models and in human AD cohorts could help scrutinize this aspect in the future. Finally, our study identified *Slc22a17* as a potential drug target to design therapies for interventions that will aim to enhance neurogenesis in AD.

Our study is strong in terms of providing a link between enhanced neurogenesis and reduced AD pathology burden. Additionally, we provide an evolutionarily conserved mechanism for NGFR signalling (from zebrafish to mammals) in regulating neurogenesis-related pathological alterations. We provide evidence for the antagonistic effects of NGFR signalling on LCN2 and reactive gliosis using two mammalian species – human and mouse. We utilized transcriptomics, spatial proteomics, *in vivo* functional knockdown studies, cell labeling and tracing as sensitive tools and identified a potentially druggable receptor, Slc22a17. Our study also has some limitations. We performed transient knockdowns *in vivo*, and transgenic animal models can be used in the future for measuring the sustained effects. Additionally, in the future, Tau pathology mouse models could be used to determine the long-term effects of neurogenesis and NGFR signalling on Tauopathies. Finally, we used a ubiquitous promoter for the viral gene expression because we sought for tracing the transduced cells for long-term. Therefore, astroglia-specific promoters would not be feasible. Our initial transduction mainly labels astrocytes, but we cannot exclude that the nearby cells could also be transduced. Based on our observations, this ratio is rather low and would not affect our conclusions drawn, yet in the future, cre-lox based cell specific recombination strategies could be employed for cell-specific recombination.

## Conclusions

We discovered that an autocrine molecular mechanism - Lipocalin-2 (Lcn2) / Solute carrier protein 22a17 (Slc22a17) axis regulates the neuroregenerative activity of astroglia by controlling the neurogenic versus reactive states. Ngfr-induced neurogenesis is concomitant to reduced amyloid pathology and Tau phosphorylation in mice. We found that in humans, LCN2 is elevated in AD and primary age-related Tauopathy patient brains, suggesting a relationship between reduced neurogenic capacity and the etiology of these diseases. By using the single cell sequencing datasets, we also show that in fetal human brains, 3D brain organoids and human AD patient brains, *NGFR* expression defines neurogenic capacity that declines with age. Comparison of our findings to human transcriptomics datasets in AD cohorts indicated candidate genes that might play a role in neurogenic switch mechanism in mammalian astroglia. We also propose that evolutionary determinants of neuroregenerative potential can be identified by cross-species comparison, and engineered induction of regenerative programs could help design novel therapeutic routes in AD in humans. Our study identified Slc22a17 as a potential drug target to design therapies for interventions to enhance neurogenesis in Alzheimer’s disease and proposed additional candidate genes that may function in the NGFR/p75NTR pathway that are poised for further validations.

## Materials and Methods

### Animal Maintenance

Mice were kept under pathogen-free conditions on a strict 12 hour alternating light and dark cycle with standard mouse food (chow) and water *ad libitum*. They were group-housed in standard ventilated cages prior to the experiments and were kept in individual cages after surgical procedures. Fixed gender mice, aged between 52-56 weeks were used for the experiments unless stated otherwise. B6.Cg-Tg(APP695)3DboTG(PSEN1dE) mice were obtained from Jackson Laboratories (Bar Harbor, ME, USA) and were maintained as a heterozygous breeding colony.

### Lentiviral Construct and production

In this study, we have designed and used p6NST90-hUb-mNgfr-T2A-mCherry-Lv16 (Lv16) and p6NST90-hUb-mCherry-Lv13 (Lv13) for lentiviral production (Fig. S1). These lentiviral plasmids were constructed using a second-generation HIV based lentiviral system, comprising of 3 plasmids, a) p6NST90 - transfer vector plasmid that contains the gene of interest (mNgfr and mCherry) ^62,63^; b) pCD/NL-BH - packaging plasmid that contains the Gag, Pol, Rev and Tat genes; and c) pczVSV-Gwt - envelope plasmid that encodes the VSV-G protein. Mouse *Ngfr* under the mammalian promoter from the human ubiquitin C gene and mCherry, separated by T2A, were cloned in the backbone of the HIV transfer vector to generate the Lv16 construct. Similarly, Lv13 was constructed without mouse *Ngfr* as a control vector (see table S1 for primer details).

To produce viral particles, we co-transfected the packaging cells (HEK293T cells) with the vectors pczVSV-Gwt and pCD/NL-BH and the transfer vector p6NST90, in a 10 cm dish with 8 ml of DMEM (10% heat-inactivated FBS, 1% Pen/Strep), 18-21 dishes per condition. 5 μg of each plasmid was mixed in 1 ml prewarmed blank DMEM (without FBS and pen/strep) and combined with 1 ml of blank DMEM containing 45 μl of polyethylenimine (PEI) before adding to the HEK293T cells. At 48 h post-transfection, the medium from the transduced cells was collected, filtered, and concentrated by several rounds of ultracentrifugation. The presence of mCherry expression was verified by fluorescent microscopy.

### Primary human astrocytes and mouse neural stem/progenitor cells culture

Primary human astrocytes (pHA) were purchased from ScienCell Research Laboratory (SRL, Catalog Number 1800) at passage one. Using complete astrocyte medium (SRL, Catalog Number 1801) supplemented with 5% fetal bovine serum (SRL, Catalog Number 0010), 1% astrocyte growth supplement (SRL, Catalog Number 1852), and 1% penicillin/streptomycin solution (SRL, Catalog Number 0503); pHA were sub-cultured to obtain passage two pHAs which were used for all experiments. Mouse neural stem/progenitor cells (NSPCs) were isolated from the dentate gyri of 3-month-old WT mice as previously described ^20,64,65^. Briefly, DGs were dissected out of the hemispheres of the freshly isolated adult mice brains on ice in PBS containing Pen/Strep. Using a scalpel, tissues were minced a little followed by enzymatic digestion with the Neural Tissue Dissociation kit from Miltenyi Biotec as per the manufacturer’s instructions. Following the dissociation, the cell suspensions were cultured in PDL/Laminin coated 25 cm^2^ flask using Neurobasal medium (GIBCO, Life Technologies), supplemented with 2% B27 (Invitrogen), 1× GlutaMAX (Life Technologies), 50 units/ml penicillin/streptomycin, 20 ng/ml EGF (Peprotech, AF-100-15), and 20 ng/ml FGF (Peprotech, AF-100-15). Cells were kept in an incubator with a 5% CO2/95% air atmosphere at 37°C and media was exchanged every 48 h. Only passages 7-10 were used during the experiments.

### Biohybrid-hydrogels 3D culture and lipocalin-2 treatment

Primary human astrocytes were encapsulated in biohybrid-hydrogels as previously described ^66^. 10-μl hydrogels were created with a concentration on 4 million cells per ml (n=3) and were treated with 200 ng/ml lipocalin-2 or PBS in a complete astrocyte medium for one week. After two weeks of culture, hydrogels were fixed using 4% PFA and were stained for GFAP markers.

### Generation of Lv13 and Lv16-transduced human astrocytes

Lv16-p6NST90-hUb-mNgfr-T2A-mCherry and Lv13-p6NST90-hUb-mCherry human astrocytes cell lines were generated using pHA pf passage 3 which were transduced with respective viral vectors. For transduction, pHA were seeded in a 24-well plate and were cultured until cells reached 70-80% confluency using a complete astrocyte medium. The respective viruses were then added to each well and cells are incubated for 24 hours at 37°C in an incubator with a 5% CO2/95% air atmosphere. After 24 hours, the media is exchanged, and cells are allowed to grow until they were 90% confluence. Cells supernatant was checked for viral load after 72 hours of infection and virus particle-free cells were passaged to a bigger vessel. Cells from overall passages 6 were used for all experiments.

### Slc22A17 and control morpholino of Lv13 and Lv16 mouse neural stem/progenitor cell lines

Mouse NSPCs from passage 7 were used to generate the Lv16-p6NST90-hUb-mNgfr-T2A-mCherry and Lv13-p6NST90-hUb-mCherry mouse NSPCs cell line. Mouse NSPCs were cultured in a PDL/Laminin coated 25 cm^2^ flask at 10,000 cells per cm^2^ concentration using Neurobasal plus medium (GIBCO, Life Technologies), supplemented with 2% B27 (Invitrogen), 1x GlutaMAX (Life Technologies), 50 units/ml penicillin/streptomycin, 20 ng/ml EGF (Peprotech, AF-100-15), and 20 ng/ml FGF (Peprotech, AF-100-15). Lv16 and Lv13 virus were then added to each flask and cells are incubated for 24 hours at 37°C in an incubator with a 5% CO2/95% air atmosphere. Media was changed after 24 hours and cells are allowed to recover and grow until they were 70-80% confluence (approx. 4 days, media change at every 48 hours). 5uM of control and Slc22A17 morpholino oligos were added to respective cell line flasks and were incubated for 24 hours. Cells were washed after 24 hours with fresh media.

### Stereotaxic injections of lentiviral vectors, lipocalin-2, and neural stem cells

The stereotaxic injection procedure was carried out as previously established protocol ^67^. Briefly, during the entire surgery, the mice were anesthetized using a mix of oxygen and isoflurane (49:1) (Baxter – HDG9623) flow and placed on a pre-warmed heat-pad to prevent hypothermia. The head was immobilized with the help of ear bars and the eyes were with a protective ointment to avoid cornea dehydration. To minimize any possible pain after the surgery, an analgesic was subcutaneously injected prior to the procedure. The hippocampal injection coordinates were ± 1.6 mm mediolateral, −1.9 mm anterior-posterior, and −1.9 mm dorsoventral from the Bregma, where the virus was dispensed at 200 nl/min speed. After the injection, the capillary was slowly retracted, followed by the release of the ear bars and stitching of the injection site.

For the viral injections of p6NST90-hUb-mNgfr-T2A-mCherry-Lv16 (Lv16) or p6NST90-hUb-mCherry-Lv13 (Lv13), 1 μl of each virus was injected into either hemisphere. For the Lcn2 injection experiment, 1 μl of 10ng/μl Lcn2 solution was injected into the right hemisphere and 1 μl vehicle solution into the left. For the injection of morpholino-treated cells, after 24 hours of morpholino treatment cells were lifted using Accutase (Gibco, A11105-01), immediately followed by a cranial injection to avoid keeping them on ice for longer than 1 hour. Each hemisphere of WT mice was delivered with 1×10^5^ cells/μl of cell suspension at a speed of 200 nl/min using the Nano-injector system. At 3 days post-injection, mice were sacrificed either via cervical dislocation followed by hippocampi extraction or via transcardial perfusion to isolate the whole brain for further processing.

### BrdU labeling and tissue preparation

To label proliferating cells, mice were injected intraperitoneally (IP) with BrdU (50 mg/kg) 3 times on the day of cranial injection, 9 hours apart, followed by an IP every 24 hours until the end of experiment. Mice were sacrificed by an overdose of Ketamine/Xylazin (0.25 mL per 25 g of body weight) mixture and perfused transcardially with NaCl (0.9% w/v) followed by 4% paraformaldehyde (PFA). Brains were harvested and post-fixed in 4% PFA at 4°C overnight. For cryopreservation of the fixed tissue, brains were transferred to a 30% sucrose solution for 2–3 days. Coronal sections with a thickness of 40 mm were cut using a sliding microtome (Leica SM2010) cooled with dry ice. Free floating sections were collected in 6 consecutive series and stored in cryoprotection solution (CPS; 25% ethylene glycol, 25% glycerol in 0.1 M phosphate buffer pH 7.4) at −20°C. Every sixth section of each brain was pooled in one series for immunohistochemistry.

### Fluorescence immunohistochemistry

Mouse: Prior to immunohistochemistry, the free-floating sections were washed in PBS 3 times, blocked in 10% Donkey or Goat Serum, 0.3% Tween 20, 1x PBS solution for one hour at room temperature. In the case of Dcx and BrdU staining, antigen retrieval was performed in 2 N HCl for 30 min at 37°C followed by 3x washing in PBS. Primary antibodies (4G8, Aldoc, BDNF, BrdU, DCX, GFAP, Hopx, LCN2, mCherry, Mcm7, NGF, NGFR, proBDNF, pTau-S199, or Slc22A17) were diluted in PBS together with 3% Goat/donkey serum and 0.3% Tween 20 and incubated overnight at 4°C with the sections. This was followed by washing with PBS 3 times within an hour and incubating for four hours at room temperature with the correct secondary antibody conjugated with a desired fluorophore. After short wash samples are then incubated in 4,6-diamidino-2-phenylindole (DAPI) diluted in PBS for 15 minutes. Additional steps of washing were done and samples are mounted on the charged glass slides. After mounting slides are left to dry and covered with a coverslip using Aqua-Poly/Mount (Polysciences Europe GmbH). Human: paraffin-embedded tissue section blanks (20-micron thickness) were obtained from New York Brain Bank. Immunohistochemistry was performed as described ^68^.

Biohybrid-hydrogels: fixed hydrogels were blocked and permeabilized using 10% Goat Serum, 0.3% Tween 20, and 1x PBS solution for one hour at room temperature, followed by primary and secondary antibody treatments (in 3% Goat serum 0.3% Tween 20, and 1x PBS). See table S1 for antibody information, table S2 for information on the human brain samples.

### Imaging and Quantifications

Spinning Disc Zeiss Axio Observer.Z1 microscope (Oberkochen, Germany) equipped with ZEN software (version blue edition, v3.2, company, Carl Zeiss, Jena, Germany) was used to acquire the fluorescence images of the 40-μm thick mouse brain tissue sections (along with the complete ventral dorsal extent of the DG at 20x magnification) and the biohybrid hydrogels (5 images per hydrogel at 10x magnification, image acquisition dimensions were 704, 999, 300 μm, respectively, with a step size of 2 μm). Human brain sections were imaged with Zeiss LSM 800 scanning confocal microscopes with Airyscan super-resolution module (Oberkochen, Germany). Images were analyzed using ZEN (v3.2, Carl Zeiss, Jena, Germany) or ImageJ (v1.53, NIH, Bethesda, MD, USA version, company, city, country, https://imagej.nih.gov/ij/) or Arivis Vision 4D (v4) software. BrdU/GFAP, BrdU/DCX/mCherry, and BrdU/GFAP/mCherry cells were manually counted in every sixth section (240 mm apart) and were normalized to the area (measured via Arivis vision 4D software) of the granular layer of the dentate gyrus present through the ventral dorsal axis of each hemisphere (Fig. 2G, Fig. S2, S3, and S4, Data S4). In the case of morpholino-treated cells injection, BrdU/DCX/mCherry triple-positive cells were manually counted, only the sections with at least one mCherry positive cell in the granular layer were considered for analysis (Fig. 2J). Quantification of amyloid β plaques (4G8 staining) and phosphorylated Tau-S199 was done via intensity measurement (using Zen v3.2 software), the total intensity was normalized to the area dentate gyrus with proper background noise correction (Fig. 3F, 3H, Data S4). Volume analysis of the hydrogel encapsulated GFAP cells and human brain sample images for Lcn2 expression were performed in an automated manner using the Arivis vision 4D pipeline.

### Single Cell Sequencing and Analyses

Mouse hippocampi were isolated as described ^20^ and cells were kept on ice until dissociated into cells which were used for the single-cell sequencing. Single cell sequencing was performed as described ^9,10,20^. Sequencing dataset is available on NCBI GEO (https://www.ncbi.nlm.nih.gov/geo/) with accession number GSE189626. We performed all analyses with R.4.0 and Seurat V4. A Seurat object was generated for each dataset, the data were normalized with NormalizeData and 500-2000 variable genes were identified. Data were scaled and nCount_RNA (nUMI) were regressed out. The first 20 PCA were determined, clusters were identified using resolution of 1 and UMAP was calculated for 2D visualization. Data scaling, cluster identification and UMAP detection was performed as described before ^10^. To identify main cell types, we used known marker genes and top marker gene of each cluster. The marker genes were identified using the Seurat function “FindAllMarkers” with only.pos = T. Then, top 20 markers of each cluster were identified and heatmaps were generated. Human organoid, fetal brain and adult brain single cell sequencing clustering, heat map generation and tSNE plots were generated as described ^9,19^.

### NanoString GeoMx digital spatial profiling (DSP)

Using DSP, we performed a multiplexed and spatially resolved profiling analysis on Lv16-p6NST90-hUb-mNgfr-T2A-mCherry and Lv13-p6NST90-hUb-mCherry injected contralateral hemispheres of WT and APP mice. DSP technology uses antibodies with UV photocleavable indexing oligos for protein profiling within the selected regions of interest (ROIs). Using the slide preparation protocol from NanoString Technologies, Inc (Seattle, WA, USA), 5-μm-thick FFPE sections of WT and APP mice brains were prepared from the viral injected site. Morphology markers for visualizing the neural cells (GFAP, AF594), immune cells (Iba1, AF647), mCherry (AF532), and nucleic acids (AF488), were applied for 1 h at room temperature prior to being loaded on the GeoMx Digital Spatial Profiler (NanoString Technologies, Inc). As shown in Fig. 5A, based on fluorescence imaging, ROIs (200–600 μm in diameter) within the DG areas were chosen for multiplex profiling. The DSP exposed each ROI to a 385-nm light to release the indexing oligos, and the photocleaved oligos were transferred into a microwell. The DSP sequencing data were processed using the GeoMx DSP analysis suite (GEOMX-B0007). Reads were normalized to signal-to-noise ratio (Fig. 3B, Data S7 and S8).

### Human Alzheimer’s disease versus control differential expression analysis

We compared the conditional-quantile-normalized^69^ gene expression levels between AD and control brains in the AMP-AD datasets ^29,31–33^ using a multiple linear regression model adjusting for the biological and technical variables (age, sex, sequencing flow-cell, RIN, tissue source, *APOE4* allele dosage). Multiple testing was adjusted using false discovery rate (FDR). The cell-intrinsic DEG (CI-DEG) analysis^34^ and the weighted gene co-expression network analysis (WGCNA) ^35,36^ were performed as previously described.

### Statistical Analyses

Data analysis was performed using Prism software (Version 8, GraphPad Software, Inc). Results were expressed as mean ± standard deviation (SD). Statistical significance was determined using t test when the experiment contained two groups, an ANOVA when comparing more than two groups, or a one-sample t test when comparing to a hypothetical value. Bonferroni and Tukey’s post hoc tests were performed and mentioned in the text wherever applicable. The level of conventional statistical significance was set at p < 0.05 and displayed visually as * p < 0.05, ** p < 0.01, and *** p < 0.001.

## Abbreviations

AD: Alzheimer’s disease
ANOVA: Analysis of variance
APP: Amyloid precursor protein
Aβ42: Amyloid-β 42
BDNF: Brain-derived neurotrophic factor
BrdU: Bromodeoxyuridine
C4B: Complement factor 4B
DAPI: 4′,6-diamidino-2-phenylindole
Dcx: Doublecortin
DEG: Differentially expressed gene
DG: Dentate gyrus
GFAP: Glial fibrillary acidic protein
Lcn2: Lipocalin-2
LV: Lentivirus
NGFR: Nerve growth factor receptor
PFKP: Phosphofructokinase, platelet
PS1: Presenilin 1
SGZ: Subgranular zone
Slc22a17: Solute carrier protein 22 a 17
tSNE: t-distributed stochastic neighbor embedding

## Declarations

### Ethical Approval

All animal experiments were performed in accordance with the applicable European regulations and approved by the responsible authority (Landesdirektion Sachsen Germany and TU Dresden-Kommission für Tierversuche) under license number TVV 87/2016. Animals were handled with extreme precaution to reduce suffering and overall animal numbers. Human brain samples were obtained from New York Brain Bank within institutional regulations of Columbia University. The analyses conducted at Mayo Clinic were approved by the appropriate Mayo Clinic Institutional Review Board.

### Availability of supporting data

All data and relevant accession numbers are available in the main text or the supplementary materials. Raw data for mouse single cell sequencing is available on GEO (accession number GSE189626). Supplementary Materials: Figs. S1 to S8, Tables S1 to S2, Data S1 to S10. Amp-AD data can be accessed via the AD Knowledge Portal. For the AMP-AD accession numbers, please use the following IDs. Bulk DEGs: ROSMAP: syn3219045 (https://www.synapse.org/#!Synapse:syn3219045); Mayo: syn5550404 (https://www.synapse.org/#!Synapse:syn5550404); MSBB: syn3159438 (https://www.synapse.org/#!Synapse:syn3159438). CI-DEG: syn22228843 (https://www.synapse.org/#!Synapse:syn22228843); WGCNA: syn5550404 (https://www.synapse.org/#!Synapse:syn5550404). The AD Knowledge Portal is a platform for accessing data, analyses and tools generated by the Accelerating Medicines Partnership (AMP-AD) Target Discovery Program and other National Institute on Aging (NIA)-supported programs to enable open-science practices and accelerate translational learning. The data, analyses and tools are shared early in the research cycle without a publication embargo on secondary use. Data is available for general research use according to the following requirements for data access and data attribution (https://adknowledgeportal.synapse.org/DataAccess/Instructions).

### Competing interests

Authors declare no competing interests.

### Funding

This work was supported by German Center for Neurodegenerative Disease (DZNE) and Columbia University Schaefer Research Scholar Award (C.K.). This work was also supported by National Institute on Aging [U01 AG046139 to N.E.T, and R01 AG061796 to N.E.T] and Alzheimer’s Association Zenith Awards (N.E.T).

### Author contributions

Conceptualization: TS, CK; Experimental materials/procedures: TS, SP, PB, ZTA, ISM, GT, CK; Experimental data acquisition/interpretation: TS, PB, CK; Single cell data analyses: TS, MIC, PB, GT, CK; Acquisition, analyses of human AMP-AD datasets: YM, XW, MA, Öİ, NE-T, Genomics cohorts and analyses: AJL, BNV, RM; Funding: CK, NE-T; Writing the manuscript: TS, PB, GT, YM, NE-T, CK; All authors contributed to editing of the manuscript. All authors read and approved the final manuscript.

## Acknowledgments

We would like to thank DRESDEN-Concept Genome Center and the deep sequencing facility, BioDIP Biopolis Dresden imaging platform and histology facility, Taub Institute for Research on Alzheimer’s Disease and the Aging Brain Imaging Platform, New York Brain Bank, and Molecular Pathology platform of the Columbia University Herbert Irving Comprehensive Cancer Center for technical help and New York Brain Bank for post-mortem human brain sections. We would like to thank Dr. Michael L. Shelanski for critical comments on the manuscript. We thank the patients and families for their participation, without whom these studies would not have been possible.

**AD Knowledge Portal: AMP-AD datasets**: The results published here are in whole or in part based on data obtained from the AMP-AD Knowledge Portal (doi:10.7303/syn2580853). **Mayo Clinic**: The Mayo RNAseq study data was led by Dr. Nilüfer Ertekin-Taner, Mayo Clinic, Jacksonville, FL as part of the multi-PI U01 AG046139 (MPIs Golde, Ertekin-Taner, Younkin, Price). Samples were provided from the following sources: The Mayo Clinic Brain Bank and Banner Sun Health Research Institute. Data collection was supported through funding by NIA grants P50 AG016574, R01 AG032990, U01 AG046139, R01 AG018023, U01 AG006576, U01 AG006786, R01 AG025711, R01 AG017216, R01 AG003949, NINDS grant R01 NS080820, CurePSP Foundation, and support from Mayo Foundation. Study data includes samples collected through the Sun Health Research Institute Brain and Body Donation Program of Sun City, Arizona. The Brain and Body Donation Program is supported by the National Institute of Neurological Disorders and Stroke (U24 NS072026 National Brain and Tissue Resource for Parkinson’s Disease and Related Disorders), the National Institute on Aging (P30 AG19610 Arizona Alzheimer’s Disease Core Center), the Arizona Department of Health Services (contract 211002, Arizona Alzheimer’s Research Center), the Arizona Biomedical Research Commission (contracts 4001, 0011, 05-901 and 1001 to the Arizona Parkinson’s Disease Consortium) and the Michael J. Fox Foundation for Parkinson’s Research. **MSBB**: These data were generated from postmortem brain tissue collected through the Mount Sinai VA Medical Center Brain Bank and were provided by Dr. Eric Schadt from Mount Sinai School of Medicine. **ROSMAP**: Study data were provided by the Rush Alzheimer’s Disease Center, Rush University Medical Center, Chicago. Data collection was supported through funding by NIA grants P30AG10161 (ROS), R01AG15819 (ROSMAP; genomics and RNAseq), R01AG17917 (MAP), R01AG30146, R01AG36042 (5hC methylation, ATACseq), RC2AG036547 (H3K9Ac), R01AG36836 (RNAseq), R01AG48015 (monocyte RNAseq) RF1AG57473 (single nucleus RNAseq), U01AG32984 (genomic and whole exome sequencing), U01AG46152 (ROSMAP AMP-AD, targeted proteomics), U01AG46161(TMT proteomics), U01AG61356 (whole genome sequencing, targeted proteomics, ROSMAP AMP-AD), the Illinois Department of Public Health (ROSMAP), and the Translational Genomics Research Institute (genomic). Additional phenotypic data can be requested at www.radc.rush.edu.

---

**Fig. S1.**
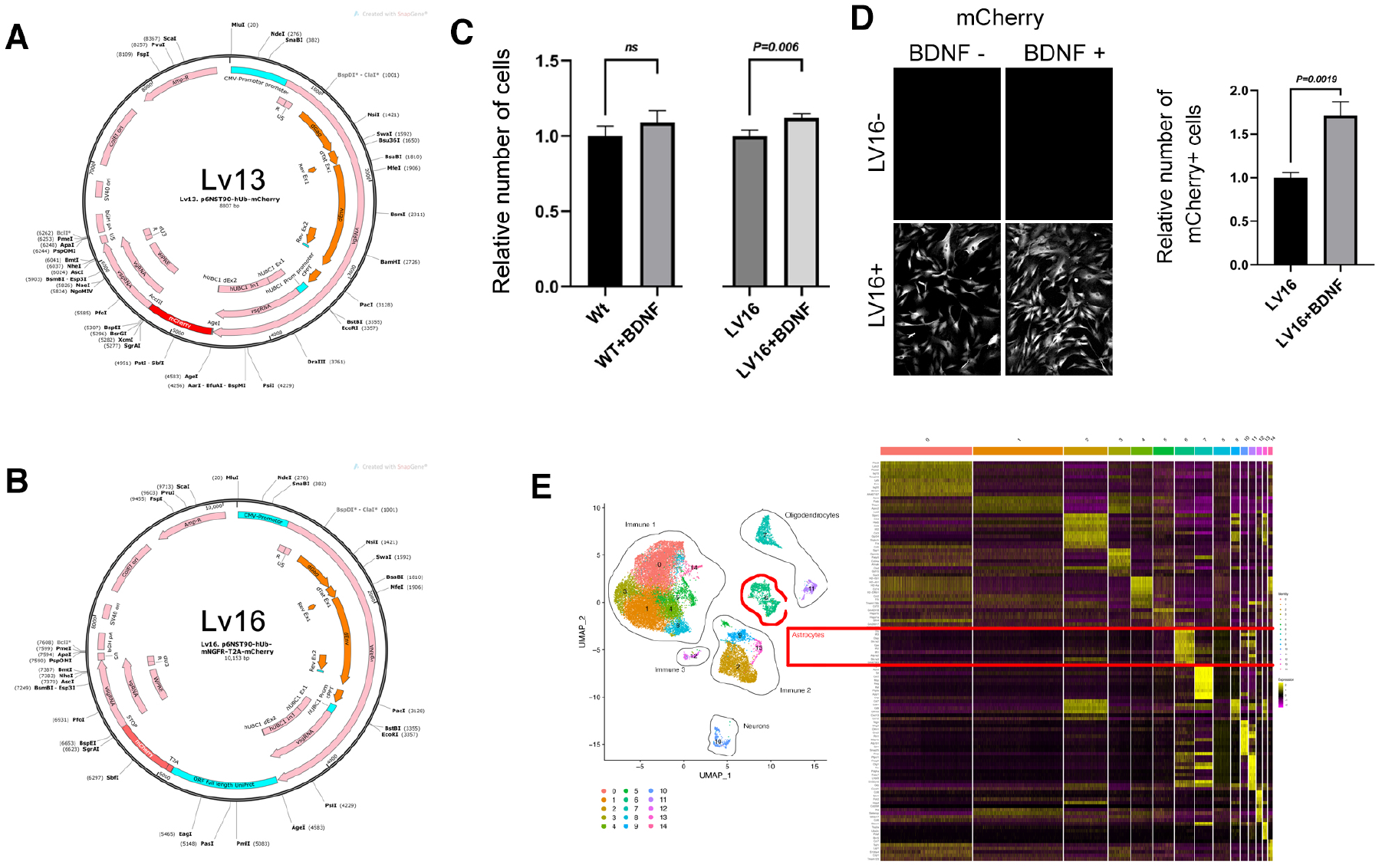
Maps of lentiviral vectors, in vitro functionality assay, single cell sequencing heat map. **(A)** Lv13: CMV promoter driven mCherry. **(B)** Lv16: CMV promoter-driven mouse Ngfr linked to mCherry with T2A self-cleaving peptide. **(C)** Graphs show relative number of cells in culture dish after equi-number seeding of cells that are either untransduced (wt) or LV16-transduced. **(D)** BDNF is added to the culture medium to activate Ngfr signalling. Only Lv16-transduced cells are responsive to BDNF. Staining image shows the mCherry -positive cells that increase in response to BDNF and Lv16. Graph indicates the number of mCherry-positive cells after equi-number seeding. **(E)** Single cell sequencing tSNE plot and heat map. Clustering of cells and their cellular identities are shown on the left. Heat map on the right shows the markers for individual cell types and their expression levels. Yellow indicates high expression. Astrocyte population marked with red rectangle.

**Fig. S2.**
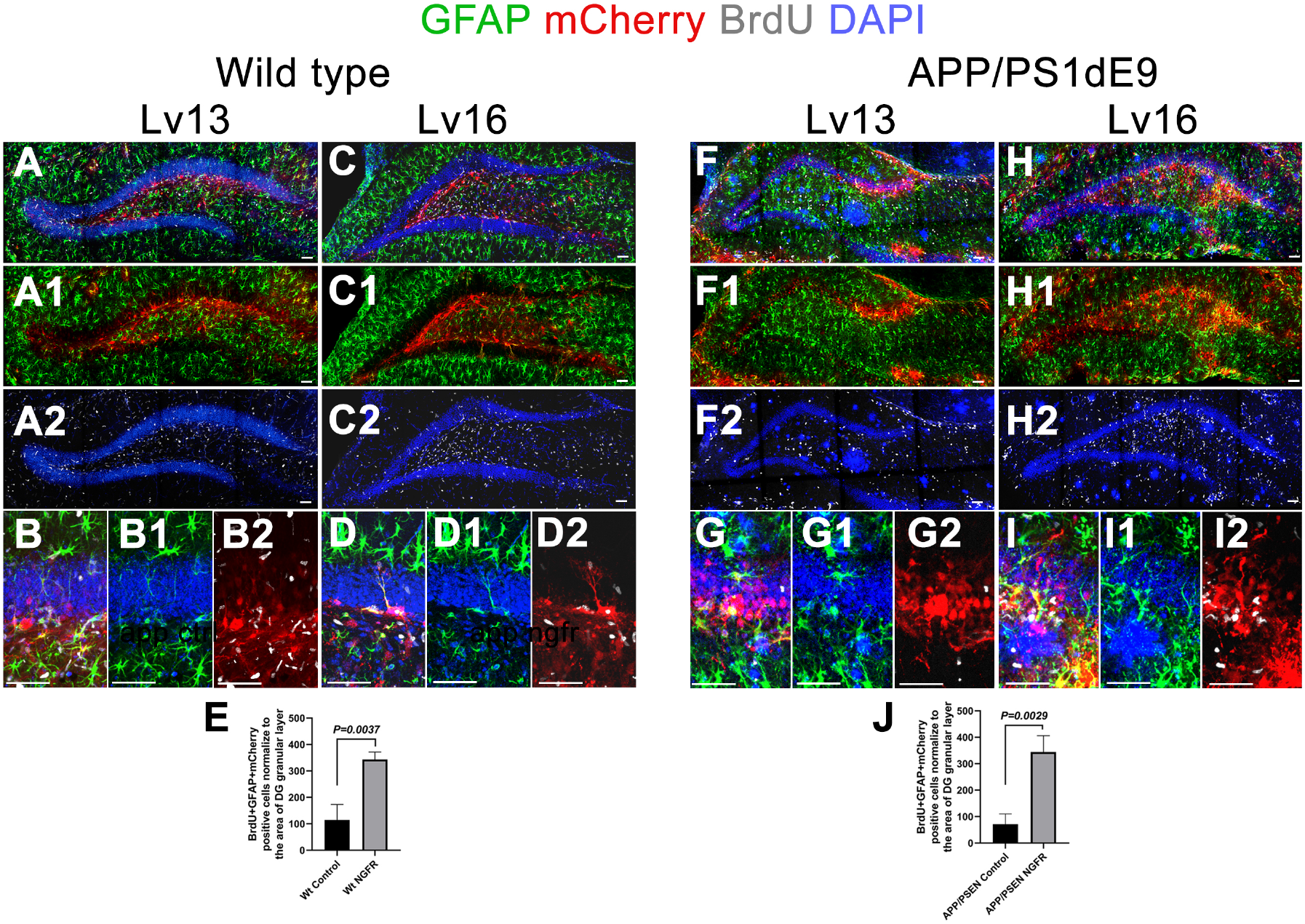
Lv16 transduction enhances astroglia proliferation in wild type and APP/PS1dE9 mouse model of Alzheimer’s disease. **(A)** GFAP, mCherry, BrdU triple immunostaining with DAPI counterstain in Lv13-transduced wild type mouse dentate gyrus. **(A1)** DAPI and BrdU omitted from A. **(A2)** GFAP and mCherry omitted from A. **(B)** Higher magnification image of subgranular zone. **(B1)** BrdU and mCherry omitted from B. **(B2)** GFAP and DAPI omitted from B. **(C)** GFAP, mCherry, BrdU triple immunostaining with DAPI counterstain in Lv16-transduced wild type mouse dentate gyrus. **(C1)** DAPI and BrdU omitted from C. **(C2)** GFAP and mCherry omitted from C. **(D)** Higher magnification image of subgranular zone. **(D1)** BrdU and mCherry omitted from D. **(D2)** GFAP and DAPI omitted from D. **(E)** Quantification graph for BrdU-mCherry-GFAP triple positive cells in Lv13 and Lv16-transduced brains. **(F)** GFAP, mCherry, BrdU triple immunostaining with DAPI counterstain in Lv13-transduced APP/PS1dE9 mouse dentate gyrus. **(F1)** DAPI and BrdU omitted from F. **(F2)** GFAP and mCherry omitted from F. **(G)** Higher magnification image of subgranular zone. **(G1)** BrdU and mCherry omitted from G. **(G2)** GFAP and DAPI omitted from G. **(H)** GFAP, mCherry, BrdU triple immunostaining with DAPI counterstain in Lv16-transduced APP/PS1dE9 mouse dentate gyrus. **(H1)** DAPI and BrdU omitted from H. **(H2)** GFAP and mCherry omitted from H. **(I)** Higher magnification image of subgranular zone. **(I1)** BrdU and mCherry omitted from I. **(I2)** GFAP and DAPI omitted from I. **(J)** Quantification graph for BrdU-mCherry-GFAP triple positive cells in Lv13 and Lv16-transduced brains. Scale bars equal 100 μm.

**Fig. S3.**
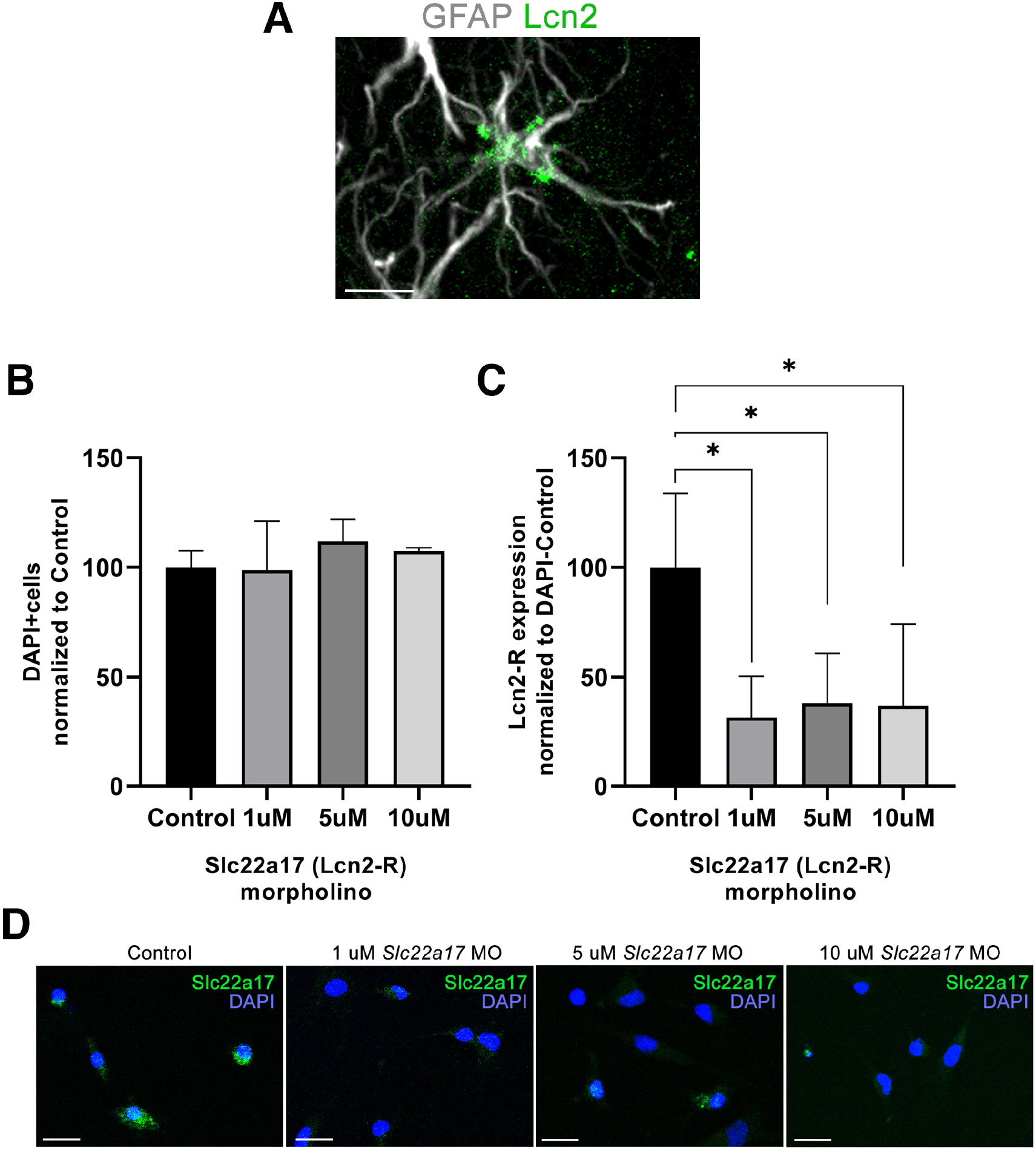
Lcn2 in astroglia and effectiveness of *Slc22a17* morpholino. **(A)** GFAP and Lcn2 double immunolabeling in mouse hippocampus. Scale bars equal 25 μm. **(B)** In vitro cell viability counts in astroglia that are untransfected or transfected with three doses of *Slc22a17* vivo morpholino (1-, 5- and 10 μM). **(C)** Slc22a17 (Lcn2-R) protein expression (fluorescence units on immunolabeled cultures) normalized to DAPI in control and transfected astroglia. **(D)** Slc22a17 immunostaining and DAPI counterstain in control (untransfected) astroglia and astroglia transfected with 1-, 5- and 10 μM vivo morpholino. Scale bars equal 25 μm.

**Fig. S4.**
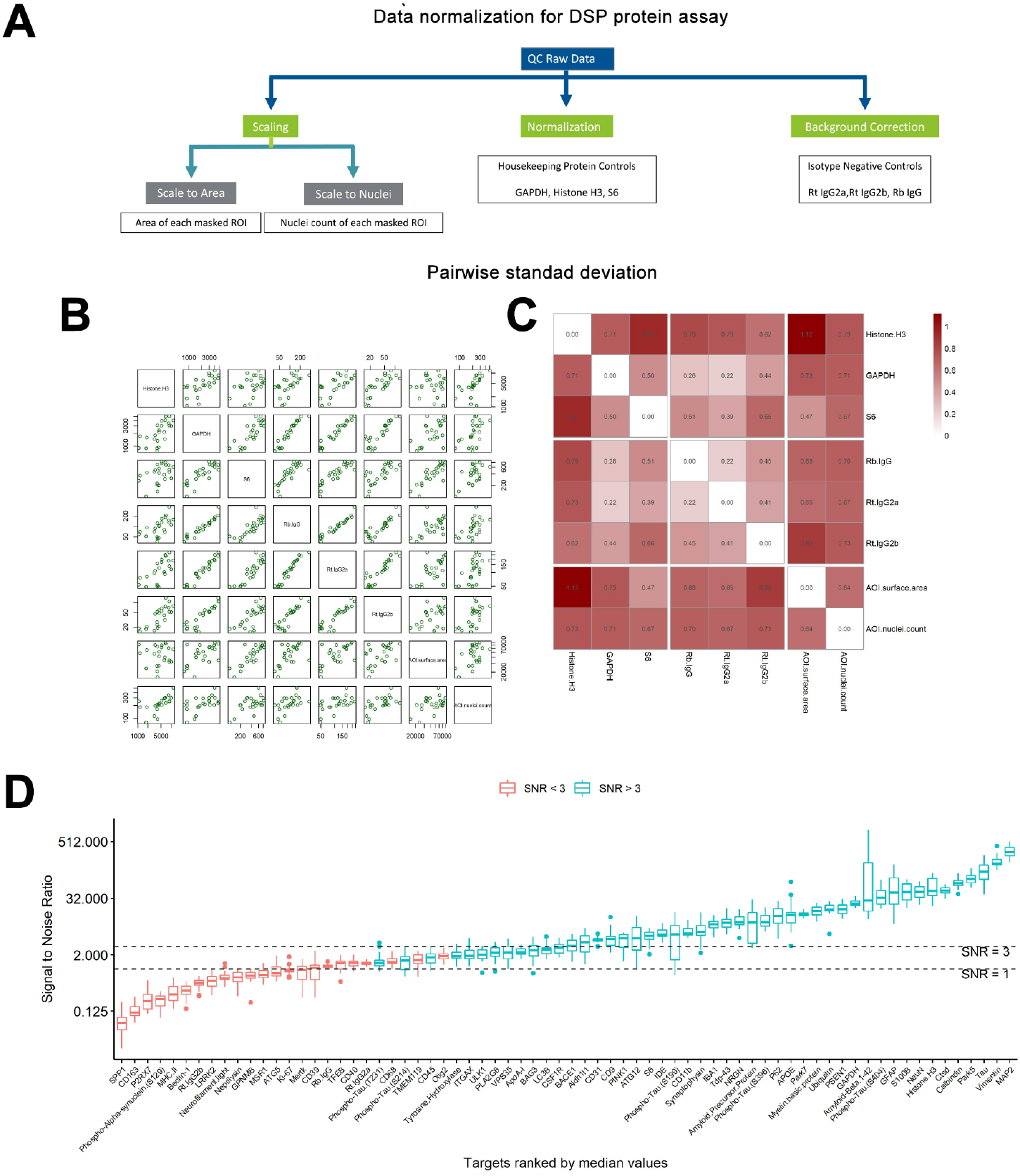
Nanostring GeoMX quality control and normalization strategy. **(A)** Data normalization options. **(B, C)** Pairwise standard deviation analysis. (B) Raw values plotted for spike-in normalization samples. (C) Standard deviation heat map for surface area, nuclei count, housekeeping genes (Histone H3, GAPDH, S6) or signal-to-noise ratio (IgG, IgG2a, IgG2b). Signal-to-noise ratio normalization has the lowest standard deviation. **(D)** Plot for targets ranked by median expression values normalized to signal-to-noise ratio.

**Fig. S5.**
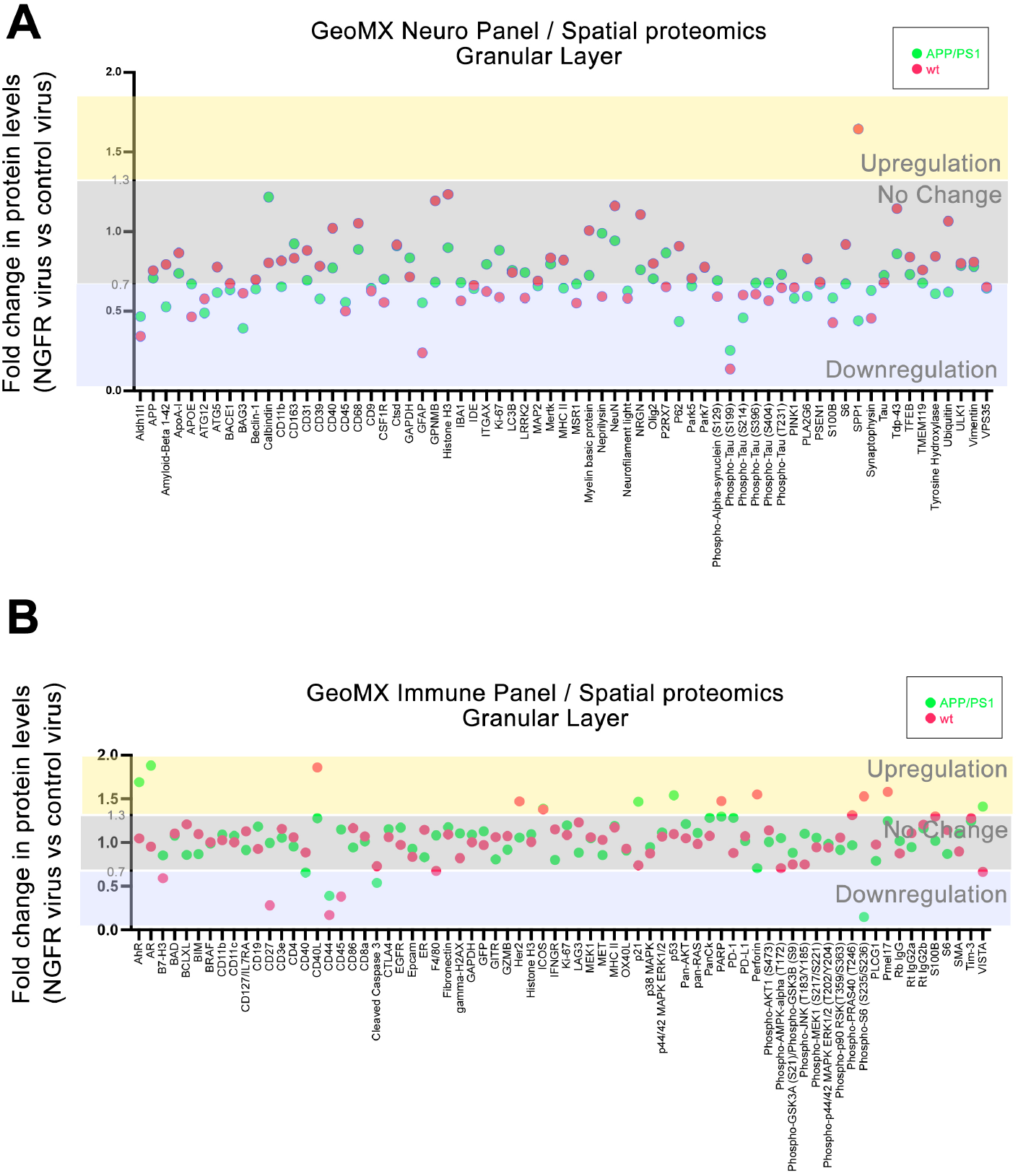
GeoMX neuro and immune panel spatial proteomics plot showing fold changes of protein targets in Ngfr-transduced brains versus control virus-transduced brains. Plots indicate targets and their fold changes in wild type animals (red dots) and APP/PS1dE9 mouse model of Alzheimer’s disease (green dots). 1.3 and 0.7 are taken as upper and lower boundaries for the no-change criterion. **(A)** Neuro panel. **(B)** Immune panel.

**Fig. S6.**
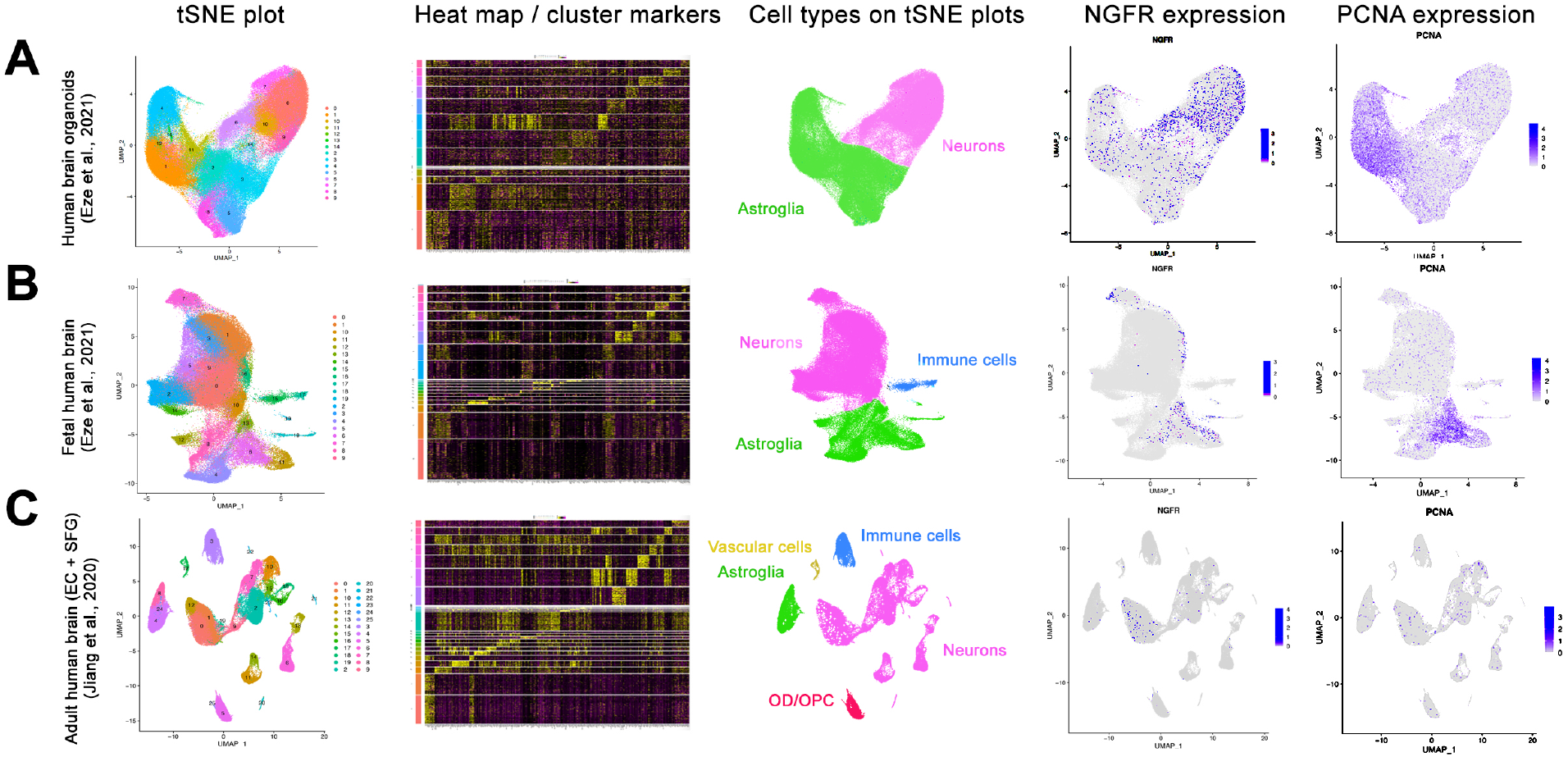
Single cell sequencing analyses of human brain organoids, fetal and adult human brains. tSNE plots for cell clustering, heat maps of top marker genes, identified cell types on colored tSNE plots, tSNE plot for *NGFR* and *PCNA* are shown for human datasets of **(A)** human brain organoids, **(B)** fetal human brain, and **(c)** adult human brain entorhinal cortex (EC) and superior frontal gyrus (SFG).

**Fig. S7.**
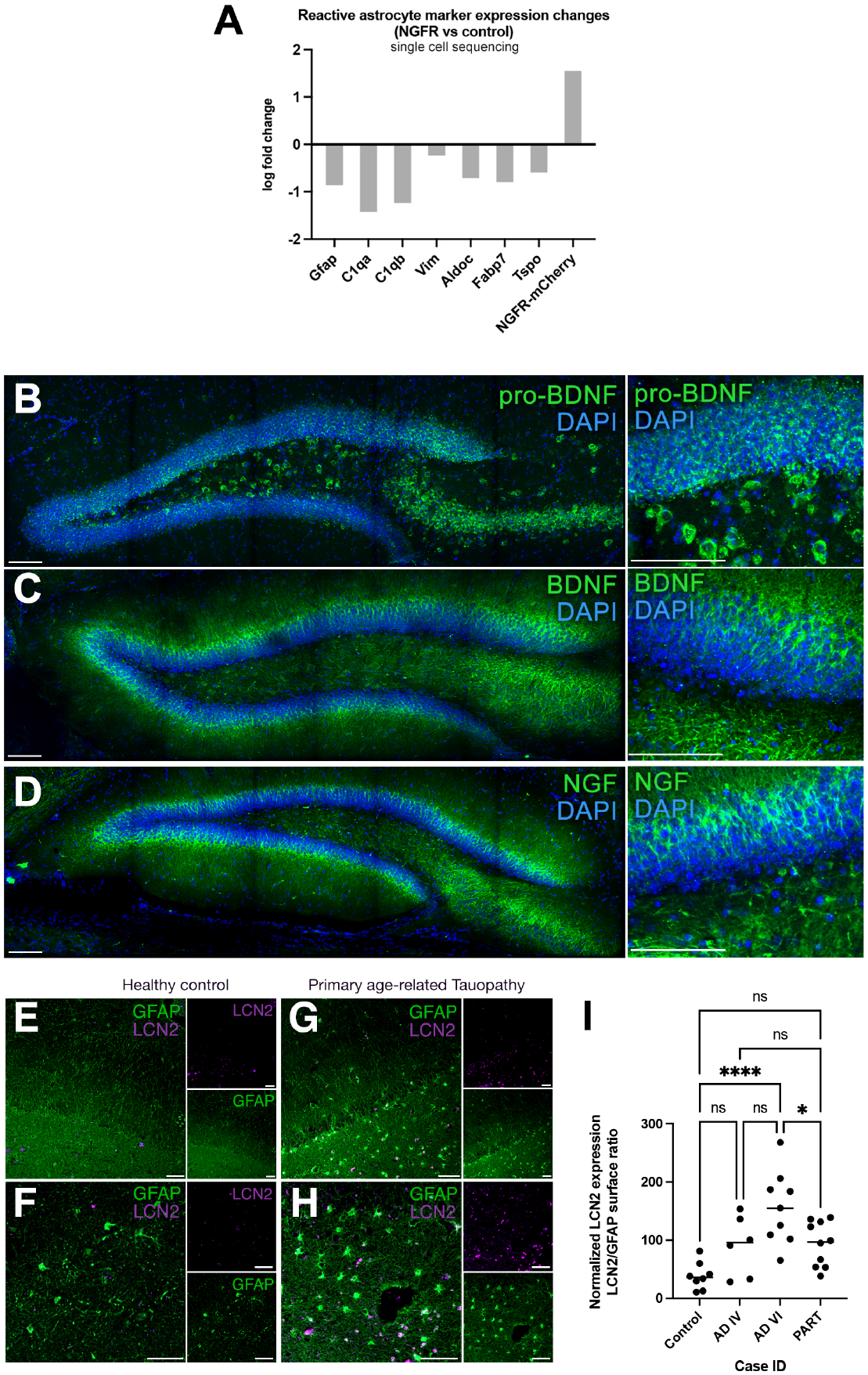
Reactive astrocyte markers in single cell transcriptomics, expression of neurotrophins in mouse dentate gyrus (DG), and LCN2 expression in primary age-related Tauopathy patients. **(A)** Expression of reactive astrocyte markers *GFAP*, *C1qa*, *C1qb*, *Vim*, *Aldoc, Fabp7*, and *Tspo* are shown. *Ngfr* expression reduces the expression of these markers. Ngfr-mCherry expression is shown for comparison. **(B-D)** Expression of pro-BDNF, BDNF and NGF in mouse dentate gyrus. Panels on the right are high magnification from left panels. **(B)** pro-BDNF, **(C)** BDNF, **(D)** NGF. BDNF and NGF are expressed in outer layers of DG, while pro-BDNF is expressed in the subgranular zone and hilus. **(E-H)** Immunohistochemical stainings for LCN2 and GFAP on hippocampal brain sections of healthy control (E, F; identical to Fig. 4) and primary age-related Tauopathy patient (G, H). Single channel insets to the right of every panel indicate individual fluorescent channels. LCN2 (top) and GFAP (bottom). **(I)** Quantification of LCN2-positive GFAP cells normalized to total GFAP cells. The samples values in Fig. 4 are also included to perform a multiple comparison. In total n = 10 human brains, n = 32 images analyzed. ****: p<0.0001. Scale bars equal 25 μm (B-D) and 50 μm (E-H).

**Fig. S8.**
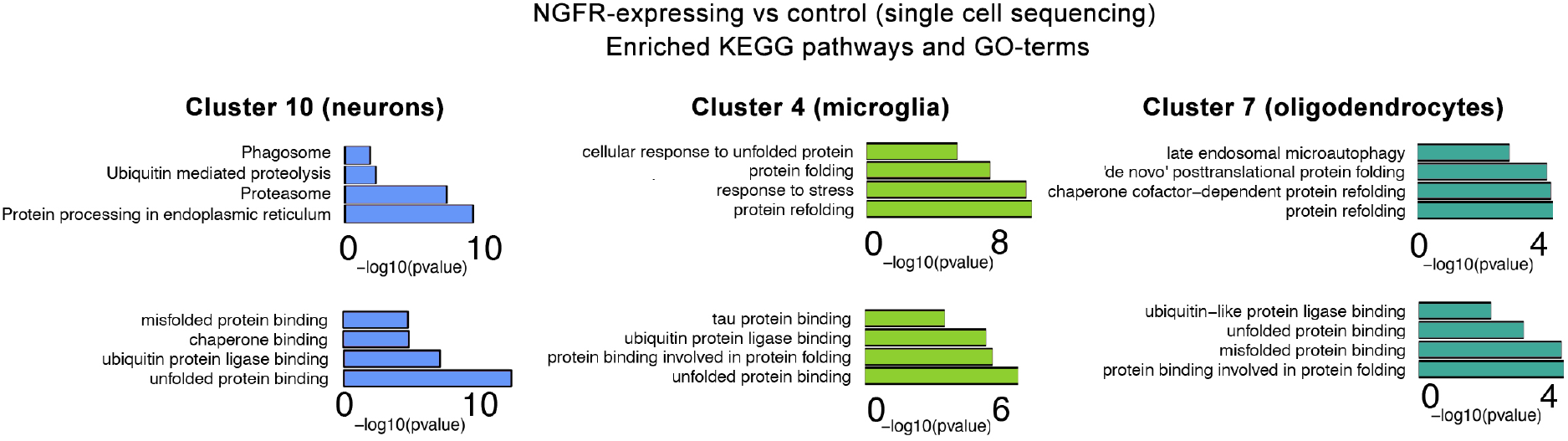
Protein clearance and misfolded protein response-related pathways are enriched upon *Ngfr* expression in non-glial cell types. Differentially expressed genes after *Ngfr* transduction upregulate the pathways related to enhanced protein quality control, toxic protein removal, misfolded protein response in non-neuronal cells. Selected GO-terms and KEGG pathways are depicted.

## Supplementary Tables and Data

**Table S1**. All materials and reagents used in this study.

**Table S2**. Demographics of human brain samples used in this study.

**Data S1**. Differentially expressed genes in cell clusters of mouse brain after transduction of *Ngfr*.

**Data S2**. KEGG pathway and GO term analyses on the genes that are downregulated in Cluster 6 (astroglia) after Lv16 transduction compared to control.

**Data S3**. KEGG pathway and GO term analyses on the genes that are upregulated in Cluster 6 (astroglia) after Lv16 transduction compared to control.

**Data S4**. All raw data that is used for statistical calculations. Tabs indicate figure panels.

**Data S5**. Differential gene expression in astrocytes that are transduced with Ngfr virus versus non-transduced.

**Data S6**. Differential gene expression in astrocytes that are transduced with Ngfr virus versus transduced with control virus.

**Data S7**. Nanostring GeoMX spatial proteomics neuro panel raw data.

**Data S8**. Nanostring GeoMX spatial proteomics immune panel raw data

**Data S9**. Comparison of Data S6 to human ROSMAP AD cohort and zebrafish.

**Data. S10**. Significant differential gene expression (DGE) (FDR < 0.05) in the AMP-AD datasets; CellCODE, PSEA, WLC analyses for cell intrinsic DGE from 3 different datasets (Mayo, MSSM, ROSMAP); WGCNA networks constructed from Mayo CER and TCX datasets.

## Notes

### Competing Interest Statement

The authors have declared no competing interest.

### Summary of Updates

New version with new data and authors

## References

1 Tanaka, E. M. & Ferretti, P. Considering the evolution of regeneration in the central nervous system. Nat Rev Neurosci 10, 713–723, doi:10.1038/nrn2707 (2009).

2 Wu, G. et al. Understanding resilience. Front Behav Neurosci 7, 10, doi:10.3389/fnbeh.2013.00010 (2013).

3 Stern, Y. Cognitive reserve in ageing and Alzheimer’s disease. The Lancet. Neurology 11, 1006–1012, doi:10.1016/S1474-4422(12)70191-6 (2012).

4 Moreno-Jimenez, E. P. et al. Adult hippocampal neurogenesis is abundant in neurologically healthy subjects and drops sharply in patients with Alzheimer’s disease. Nat Med 25, 554–560, doi:10.1038/s41591-019-0375-9 (2019).

5 Terreros-Roncal, J. et al. Impact of neurodegenerative diseases on human adult hippocampal neurogenesis. Science, eabl5163, doi:10.1126/science.abl5163 (2021).

6 Cosacak, M. I., Papadimitriou, C. & Kizil, C. Regeneration, Plasticity, and Induced Molecular Programs in Adult Zebrafish Brain. Biomed Res Int 2015:769763(2015).

7 Kizil, C., Kaslin, J., Kroehne, V. & Brand, M. Adult neurogenesis and brain regeneration in zebrafish. Dev Neurobiol 72, 429–461, doi:10.1002/dneu.20918 (2012).

8 Jurisch-Yaksi, N., Yaksi, E. & Kizil, C. Radial glia in the zebrafish brain: Functional, structural, and physiological comparison with the mammalian glia. Glia, doi:10.1002/glia.23849 (2020).

9 Bhattarai, P. et al. Neuron-glia interaction through Serotonin-BDNF-NGFR axis enables regenerative neurogenesis in Alzheimer’s model of adult zebrafish brain.. PLoS Biol 18, e3000585, doi:10.1371/journal.pbio.3000585. (2020).

10 Cosacak, M. I. et al. Single-Cell Transcriptomics Analyses of Neural Stem Cell Heterogeneity and Contextual Plasticity in a Zebrafish Brain Model of Amyloid Toxicity. Cell Rep 27, 1307–1318 e1303, doi:10.1016/j.celrep.2019.03.090 (2019).

11 Bhattarai, P. et al. Modeling Amyloid-β42 Toxicity and Neurodegeneration in Adult Zebrafish Brain. Journal of Visualized Experiments 128, doi:10.3791/56014 (2017).

12 Bhattarai, P. et al. IL4/STAT6 signaling activates neural stem cell proliferation and neurogenesis upon Amyloid-β42 aggregation in adult zebrafish brain. Cell Reports 17, 941–948, doi:10.1016/j.celrep.2016.09.075 (2016).

13 Cosacak, M. I. et al. Single Cell/Nucleus Transcriptomics Comparison in Zebrafish and Humans Reveals Common and Distinct Molecular Responses to Alzheimer’s Disease. Cells 11, 1807, doi:https://doi.org/10.3390/cells11111807 (2022).

14 Turgutalp, B. et al. Discovery of Potent Cholinesterase Inhibition-Based Multi-Target-Directed Lead Compounds for Synaptoprotection in Alzheimer’s Disease. J Med Chem (2022).

15 Bhattarai, P., Turgutalp, B. & Kizil, C. Zebrafish as an Experimental and Preclinical Model for Alzheimer’s Disease. ACS Chem Neurosci 13, 2939–2941, doi:10.1021/acschemneuro.2c00583 (2022).

16 Reinhardt, L. et al. Dual Inhibition of GSK3beta and CDK5 Protects the Cytoskeleton of Neurons from Neuroinflammatory-Mediated Degeneration In Vitro and In Vivo. Stem cell reports 12, 502–517, doi:10.1016/j.stemcr.2019.01.015 (2019).

17 Kizil, C. et al. Admixture Mapping of Alzheimer’s disease in Caribbean Hispanics identifies a new locus on 22q13.1. Mol Psychiatry, doi:10.1038/s41380-022-01526-6 (2022).

18 Lee, A. J. et al. FMNL2 regulates gliovascular interactions and is associated with vascular risk factors and cerebrovascular pathology in Alzheimer’s disease. Acta neuropathologica, doi:10.1007/s00401-022-02431-6 (2022).

19 Cosacak, M. I., Bhattarai, P. & Kizil, C. Alzheimer’s disease, neural stem cells and neurogenesis: cellular phase at single-cell level. Neural Reg Res 15, 824–827, doi:10.4103/1673-5374.268896 (2020).

20 Mashkaryan, V. et al. Type 1 Interleukin-4 signaling obliterates mouse astroglia in vivo but not in vitro Frontiers in Cell and Developmental Biology 8, 114, doi:10.3389/fcell.2020.00114 (2020).

21 Betten, R. et al. Tonicity inversely modulates lipocalin-2 (Lcn2/24p3/NGAL) receptor (SLC22A17) and Lcn2 expression via Wnt/beta-catenin signaling in renal inner medullary collecting duct cells: implications for cell fate and bacterial infection. Cell Commun Signal 16, 74, doi:10.1186/s12964-018-0285-3 (2018).

22 Ishii, A. et al. Obesity-promoting and anti-thermogenic effects of neutrophil gelatinase-associated lipocalin in mice. Sci Rep 7, 15501, doi:10.1038/s41598-017-15825-4 (2017).

23 Arber, C. et al. Familial Alzheimer’s Disease Mutations in PSEN1 Lead to Premature Human Stem Cell Neurogenesis. Cell Rep 34, 108615, doi:10.1016/j.celrep.2020.108615 (2021).

24 Choi, S. H. & Tanzi, R. E. Is Alzheimer’s Disease a Neurogenesis Disorder? Cell stem cell 25, 7–8, doi:10.1016/j.stem.2019.06.001 (2019).

25 Choi, S. H. et al. Combined adult neurogenesis and BDNF mimic exercise effects on cognition in an Alzheimer’s mouse model. Science 361, doi:10.1126/science.aan8821 (2018).

26 Kizil, C. & Bhattarai, P. Is Alzheimer’s Also a Stem Cell Disease? - The Zebrafish Perspective. Front Cell Dev Biol 6, 159, doi:10.3389/fcell.2018.00159 (2018).

27 Eze, U. C., Bhaduri, A., Haeussler, M., Nowakowski, T. J. & Kriegstein, A. R. Single-cell atlas of early human brain development highlights heterogeneity of human neuroepithelial cells and early radial glia. Nat Neurosci 24, 584–594, doi:10.1038/s41593-020-00794-1 (2021).

28 Jiang, J., Wang, C., Qi, R., Fu, H. & Ma, Q. scREAD: A Single-Cell RNA-Seq Database for Alzheimer’s Disease. iScience 23, 101769, doi:10.1016/j.isci.2020.101769 (2020).

29 De Jager, P. L. et al. A multi-omic atlas of the human frontal cortex for aging and Alzheimer’s disease research. Sci Data 5, 180142, doi:10.1038/sdata.2018.142 (2018).

30 Bennett, D. A. et al. Religious Orders Study and Rush Memory and Aging Project. J Alzheimers Dis 64, S161–S189, doi:10.3233/JAD-179939 (2018).

31 Hodes, R. J. & Buckholtz, N. Accelerating Medicines Partnership: Alzheimer’s Disease (AMP-AD) Knowledge Portal Aids Alzheimer’s Drug Discovery through Open Data Sharing. Expert Opin Ther Targets 20, 389–391, doi:10.1517/14728222.2016.1135132 (2016).

32 Allen, M. et al. Human whole genome genotype and transcriptome data for Alzheimer’s and other neurodegenerative diseases. Sci Data 3, 160089, doi:10.1038/sdata.2016.89 (2016).

33 Wang, M. et al. The Mount Sinai cohort of large-scale genomic, transcriptomic and proteomic data in Alzheimer’s disease. Sci Data 5, 180185, doi:10.1038/sdata.2018.185 (2018).

34 Wang, X. et al. Deciphering cellular transcriptional alterations in Alzheimer’s disease brains. Mol Neurodegener 15, 38, doi:10.1186/s13024-020-00392-6 (2020).

35 Langfelder, P. & Horvath, S. WGCNA: an R package for weighted correlation network analysis. BMC Bioinformatics 9, 559, doi:10.1186/1471-2105-9-559 (2008).

36 Conway, O. J. et al. ABI3 and PLCG2 missense variants as risk factors for neurodegenerative diseases in Caucasians and African Americans. Mol Neurodegener 13, 53, doi:10.1186/s13024-018-0289-x (2018).

37 Allen, M. et al. Conserved brain myelination networks are altered in Alzheimer’s and other neurodegenerative diseases. Alzheimers Dement 14, 352–366, doi:10.1016/j.jalz.2017.09.012 (2018).

38 Yao, X. Q. et al. p75NTR ectodomain is a physiological neuroprotective molecule against amyloid-beta toxicity in the brain of Alzheimer’s disease. Mol Psychiatry 20, 1301–1310, doi:10.1038/mp.2015.49 (2015).

39 Huang, E. J. & Reichardt, L. F. Trk receptors: roles in neuronal signal transduction. Annu Rev Biochem 72, 609–642, doi:10.1146/annurev.biochem.72.121801.161629 (2003).

40 Blurton-Jones, M. et al. Neural stem cells improve cognition via BDNF in a transgenic model of Alzheimer disease. Proceedings of the National Academy of Sciences of the United States of America 106, 13594–13599, doi:10.1073/pnas.0901402106 (2009).

41 Nykjaer, A., Willnow, T. E. & Petersen, C. M. p75NTR--live or let die. Curr Opin Neurobiol 15, 49–57, doi:10.1016/j.conb.2005.01.004 (2005).

42 De Vincenti, A. P., Rios, A. S., Paratcha, G. & Ledda, F. Mechanisms That Modulate and Diversify BDNF Functions: Implications for Hippocampal Synaptic Plasticity. Front Cell Neurosci 13, 135, doi:10.3389/fncel.2019.00135 (2019).

43 Lee, S. et al. Lipocalin-2 is an autocrine mediator of reactive astrocytosis. J Neurosci 29, 234–249, doi:10.1523/JNEUROSCI.5273-08.2009 (2009).

44 Zhao, N. et al. Lipocalin-2 may produce damaging effect after cerebral ischemia by inducing astrocytes classical activation. Journal of neuroinflammation 16, 168, doi:10.1186/s12974-019-1556-7 (2019).

45 Llorens, F. et al. Cerebrospinal fluid lipocalin 2 as a novel biomarker for the differential diagnosis of vascular dementia. Nat Commun 11, 619, doi:10.1038/s41467-020-14373-2 (2020).

46 Yamada, Y. et al. Lipocalin 2 attenuates iron-related oxidative stress and prolongs the survival of ovarian clear cell carcinoma cells by up-regulating the CD44 variant. Free Radic Res 50, 414–425, doi:10.3109/10715762.2015.1134795 (2016).

47 Young, K. M., Merson, T. D., Sotthibundhu, A., Coulson, E. J. & Bartlett, P. F. p75 neurotrophin receptor expression defines a population of BDNF-responsive neurogenic precursor cells. J Neurosci 27, 5146–5155, doi:10.1523/JNEUROSCI.0654-07.2007 (2007).

48 Angelastro, J. M. et al. Identification of diverse nerve growth factor-regulated genes by serial analysis of gene expression (SAGE) profiling. Proc Natl Acad Sci U S A 97, 10424–10429, doi:10.1073/pnas.97.19.10424 (2000).

49 Maurage, C. A., Sergeant, N., Ruchoux, M. M., Hauw, J. J. & Delacourte, A. Phosphorylated serine 199 of microtubule-associated protein tau is a neuronal epitope abundantly expressed in youth and an early marker of tau pathology. Acta neuropathologica 105, 89–97, doi:10.1007/s00401-002-0608-7 (2003).

50 Ermert, D. & Blom, A. M. C4b-binding protein: The good, the bad and the deadly. Novel functions of an old friend. Immunol Lett 169, 82–92, doi:10.1016/j.imlet.2015.11.014 (2016).

51 Finehout, E. J., Franck, Z. & Lee, K. H. Complement protein isoforms in CSF as possible biomarkers for neurodegenerative disease. Dis Markers 21, 93–101, doi:10.1155/2005/806573 (2005).

52 Heneka, M. T. et al. Neuroinflammation in Alzheimer’s disease. The Lancet. Neurology 14, 388–405, doi:10.1016/S1474-4422(15)70016-5 (2015).

53 Hudson, L. C., Bragg, D. C., Tompkins, M. B. & Meeker, R. B. Astrocytes and microglia differentially regulate trafficking of lymphocyte subsets across brain endothelial cells. Brain Res 1058, 148–160, doi:10.1016/j.brainres.2005.07.071 (2005).

54 Naude, P. J. et al. Lipocalin 2: novel component of proinflammatory signaling in Alzheimer’s disease. FASEB J 26, 2811–2823, doi:10.1096/fj.11-202457 (2012).

55 Saito, E. R. et al. Alzheimer’s disease alters oligodendrocytic glycolytic and ketolytic gene expression. Alzheimers Dement 17, 1474–1486, doi:10.1002/alz.12310 (2021).

56 March-Diaz, R. et al. Hypoxia compromises the mitochondrial metabolism of Alzheimer’s disease microglia via HIF1. Nature Aging 1, 385–399, doi:https://doi.org/10.1038/s43587-021-00054-2 (2021).

57 Wyss-Coray, T. et al. Adult mouse astrocytes degrade amyloid-beta in vitro and in situ. Nat Med 9, 453–457, doi:10.1038/nm838 [pii] (2003).

58 Kronenberg, G. et al. Subpopulations of proliferating cells of the adult hippocampus respond differently to physiologic neurogenic stimuli. J Comp Neurol 467, 455–463, doi:10.1002/cne.10945 (2003).

59 Casse, F., Richetin, K. & Toni, N. Astrocytes’ Contribution to Adult Neurogenesis in Physiology and Alzheimer’s Disease. Front Cell Neurosci 12, 432, doi:10.3389/fncel.2018.00432 (2018).

60 Yamazaki, Y. & Kanekiyo, T. Blood-Brain Barrier Dysfunction and the Pathogenesis of Alzheimer’s Disease. Int J Mol Sci 18, doi:10.3390/ijms18091965 (2017).

61 Dzamba, D., Harantova, L., Butenko, O. & Anderova, M. Glial cells - the key elements of Alzheimer′s disease. Curr Alzheimer Res (2016).

62 Ho, Y. P., Schnabel, V., Swiersy, A., Stirnnagel, K. & Lindemann, D. A small-molecule-controlled system for efficient pseudotyping of prototype foamy virus vectors. Mol Ther 20, 1167–1176, doi:10.1038/mt.2012.61 (2012).

63 Stirnnagel, K. et al. Analysis of prototype foamy virus particle-host cell interaction with autofluorescent retroviral particles. Retrovirology 7, 45, doi:10.1186/1742-4690-7-45 (2010).

64 Walker, D. G., Dalsing-Hernandez, J. E., Campbell, N. A. & Lue, L.-F. Decreased expression of CD200 and CD200 receptor in Alzheimer’s disease: a potential mechanism leading to chronic inflammation. Experimental neurology 215, 5–19, doi:10.1016/j.expneurol.2008.09.003 (2009).

65 Walker, T. L. & Kempermann, G. One mouse, two cultures: isolation and culture of adult neural stem cells from the two neurogenic zones of individual mice. J Vis Exp, e51225, doi:10.3791/51225 (2014).

66 Papadimitriou, C. et al. 3D Culture Method for Alzheimer’s Disease Modeling Reveals Interleukin-4 Rescues Abeta42-Induced Loss of Human Neural Stem Cell Plasticity. Dev Cell 46, 85–101 e108, doi:10.1016/j.devcel.2018.06.005 (2018).

67 Artegiani, B., Lindemann, D. & Calegari, F. Overexpression of cdk4 and cyclinD1 triggers greater expansion of neural stem cells in the adult mouse brain. J Exp Med 208, 937–948, doi:10.1084/jem.20102167 (2011).

68 Waldvogel, H. J., Curtis, M. A., Baer, K., Rees, M. I. & Faull, R. L. Immunohistochemical staining of post-mortem adult human brain sections. Nat Protoc 1, 2719–2732, doi:10.1038/nprot.2006.354 (2006).

69 Hansen, K. D., Irizarry, R. A. & Wu, Z. Removing technical variability in RNA-seq data using conditional quantile normalization. Biostatistics 13, 204–216, doi:10.1093/biostatistics/kxr054 (2012).

